# Lateral entorhinal cortex supports behaviorally-induced hippocampal ensemble stability for reliable memory recall

**DOI:** 10.64898/2026.03.23.711409

**Authors:** Maya D. Hopkins, Pauline Rahal, Vincent Robert, Esther Kim, Jayeeta Basu

**Author notes:** Corresponding Authors: Jayeeta Basu.

## Abstract

Hippocampal pyramidal neurons function as place cells, showing location-specific activity during navigation, to form an internal spatial map of the environment. They are hypothesized to be the neural substrate of episodic memory. However, place cell receptive fields tend to drift or have poor tuning in low demand tasks, lacking operant goals such as random foraging, or in sensory context-deprived environments. Through chronic two-photon calcium imaging of hippocampal area CA1, we directly compare stability in a low versus a high demand task within the same animals over the course of learning and recall in the same environment. We find that compared to random foraging, an odor-context based navigational task stabilizes place cell representations and increases place cell quality and quantity. To investigate the circuit mechanism that may support this stability, we manipulated the activity of lateral entorhinal cortex (LEC) excitatory neurons, which provide both indirect and direct multisensory inputs about context, odor, and time to CA1. We chemogenetically suppressed activity of excitatory neurons in LEC during recall of the odor-context based navigation task and found that context discrimination is impaired at both the behavioral and neural level. With LEC silencing, mice had lower behavioral performance, less stable population activity, and greater similarity between opposing trial types. Our study finds that increasing task demand increases CA1 stability and that this stability is partially supported by LEC.

## Introduction

Behaviorally driven functional interactions between the entorhinal cortex and the hippocampus are critical for episodic memory and goal-directed spatial navigation^1–3^. Multisensory information is integrated into a spatial code that represents an animal’s environment and its specific experiences within that environment^4,5^. To be an efficient system, this hippocampal code must be both robust and flexible. For example, hippocampal neurons form spatially tuned place maps that reactivate when an animal returns to a familiar environment^6^. Conversely, place representations can “remap” and adaptively change their spatial tuning or activity rate in response to a new environment or task rule^7–14^. This spectrum of stability and flexibility serves the overall goal of accurately representing an experience. However, recent studies have found a surprising degree of remapping of place cells over time within the same environment^15–17^. While this phenomenon, termed representational drift^18^, may help keep memories labile for future updates or timestamp discontinuous experiences^19–21^, it calls into question the classical notion that hippocampal place cells support long-term spatial episodic memories. How can memories remain stable if the underlying neural representations supporting them drift with time? Work from our group^22^ and others^5,23–27^ suggests that sensory enrichment, behavioral demand, and task engagement stabilize and expand these otherwise drifting neural representations. However, the neural mechanisms that modulate this stability in a behaviorally context-dependent manner remain elusive. Here, we postulate that multisensory cortical input helps maintain the stability of place maps, thereby supporting long-term memory recall.

Hippocampal pyramidal neurons integrate spatial information from the medial entorhinal cortex (MEC)^28–31^ with contextual information about salient sensory cues such as odors, novelty, reward goals, task rules, and time from the lateral entorhinal cortex (LEC)^32–42^. Although canonically considered the major driver of spatial input, silencing MEC through lesions or genetic manipulations has only a modest impact on place cells, increasing place field width, but not abolishing them entirely^43,44^. This demonstrates that CA1 place coding is not simply transferred from MEC but supported by some parallel information streams. LEC lesions impairs rate remapping in CA3 place cells^45^. Moreover, recent studies with silencing of LEC input, demonstrate its role in learning-driven stabilization of CA3 place maps^46^ and cortical representations^47^. Furthermore, behavioral studies have begun to implicate LEC in high memory load tasks, context-dependent spatial navigation, and non-spatial olfactory associational tasks^42,48–54^. Collectively, these studies support the idea that, although traditionally considered non-spatial, LEC input could contribute to maintenance of stable place maps governed by spatio-contextual associational learning rules. However, this idea remains to be directly tested in CA1 during long-term memory recall. Here, we probed the impact of LEC on hippocampal CA1 context-dependent spatial coding and memory recall.

In this study, we addressed both the behavioral and circuit mechanisms underlying the stabilization of hippocampal place cells by testing the hypothesis that increasing context-dependent behavioral task demand stabilizes place maps, and that silencing LEC would impair this experience-driven stability during memory recall, thereby compromising behavioral performance. To directly test our hypothesis, we used two-photon calcium imaging to chronically track neurons in the same animals across two tasks of differing demand in the same physical environment: a low-demand random foraging task and a high-demand goal-oriented navigation task in which animals operantly learn an odor-context-place association rule. We chose this high-demand task, where head-fixed mice learn to distinguish between two odors to find water rewards at fixed locations on a treadmill track, as it triggers the emergence of context-selective place maps that remain stable during memory recall^22^. To establish a circuit mechanism that supports the high-demand task, we chemogenetically silenced excitatory cells in LEC, known to code for odor contexts and task rules, during memory recall while concurrently performing calcium imaging. We validated that LEC supports behavioral performance and the maintenance of stable hippocampal neural activity underlying memory-guided context-place coding.

## Results

### Mice learn tasks of both low and high demand within the same environment

To examine the stability of CA1 ensembles with increasing task demand, mice performed two tasks of varying complexity within the same physical environment. Mice were initially trained to perform a low-demand head-fixed task where they randomly foraged (RF) for water rewards on a 200 cm linear treadmill belt. On each lap, water rewards were operantly available at 3 new, randomized zones (Fig. 1A). Mice adopted the behavioral strategy of licking uniformly at all locations along the belt (Fig. 1B-1C). Following successful completion of random foraging, mice were trained on a more complex odor-cued goal-oriented learning (OCGOL) task previously developed and characterized by our group (Fig. 1A)^22^. After learning OCGOL over the course of 1-2 weeks, mice became expert performers (>80% correct trials for 2 consecutive days) and entered the recall stage of the task. Given a specific odor cue at the start of each lap, mice navigated to and collected water rewards from the appropriate reward zone and suppressed licking at the reward zone associated with the opposing odor cue (Fig. 1B-1D). At the recall stage, for a given imaging session, A and B trials were randomized; however, for simplicity, all subsequent analyses were filtered by trial type. Licks were significantly concentrated at reward zones in OCGOL, but not in RF, demonstrating that the mice used spatial cues to efficiently perform the task (Fig. 1D). The fraction of licks in the respective reward zones was not statistically different between A and B laps. Given the stable, learned reward zones for OCGOL, mice ran faster compared to their speed in RF sessions (Fig. 1E). However, we ensured that the mice ran similar numbers of total laps run for each task type, and all downstream analyses were restricted to run epochs (Fig. 1F).

**Figure 1:**
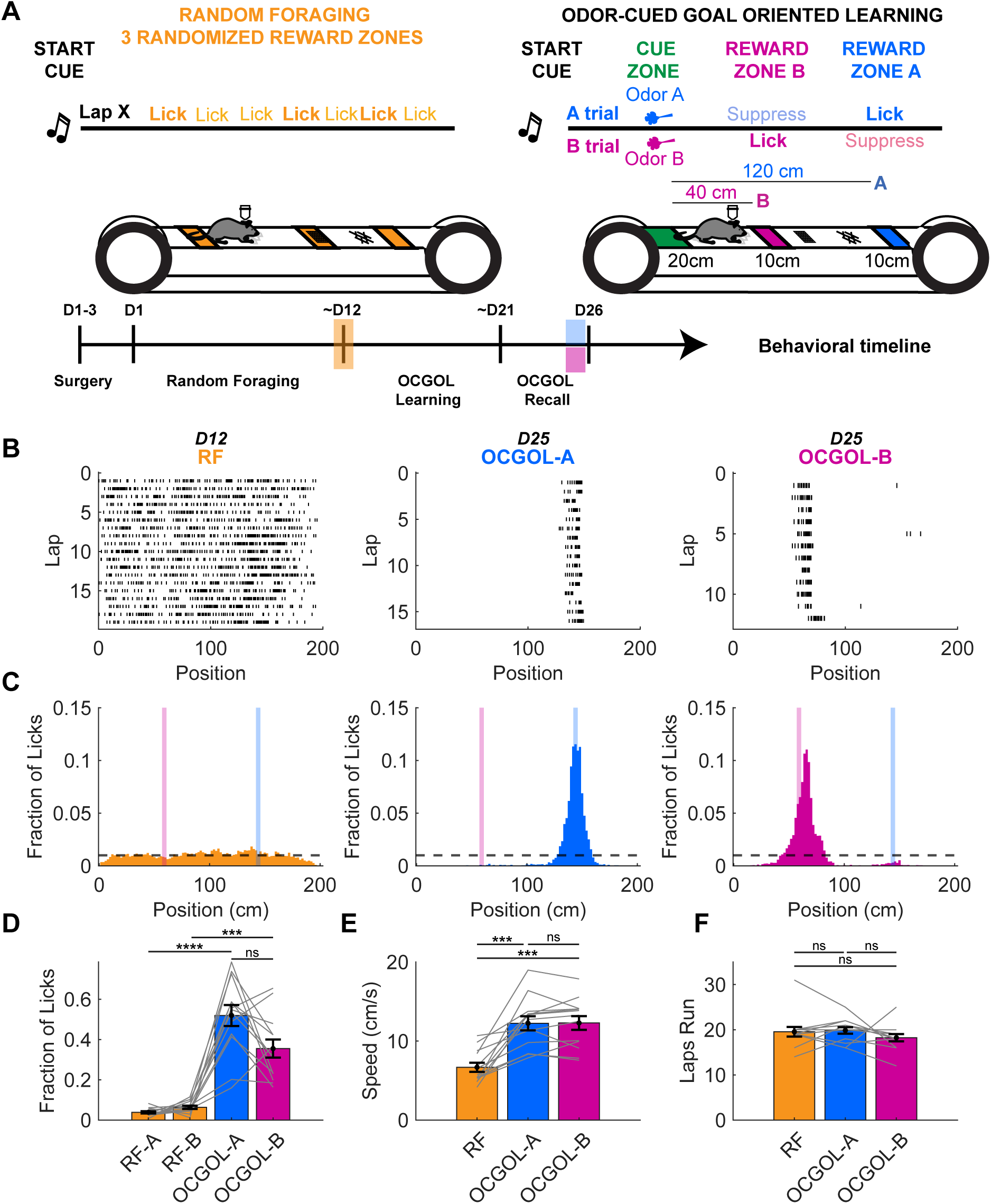
Mice learn tasks of both low and high demand within the same environment. A) Schematic of behavioral tasks: RF (left) and OCGOL (right) and behavioral timeline (bottom). B) Example lap by lap lick raster plot for a single mouse in both tasks. OCGOL is split into A (middle) and B (right) trials. C) Mean fraction of licks per 2-cm spatial bin of the treadmill belt across all mice in both tasks. OCGOL is split into A (middle, blue) and B (right, magenta) trials. Highlighted regions represent A reward zone (light blue) and B reward zone (light magenta). D) Lick fraction within the designated zone (task type-reward zone) – (one-way RM ANOVA, p<0.0001; Tukey *post-hoc* test: RF-A vs. RF-B, p=0.2634; RF-A vs. OCGOL-A, p<0.0001; RF-B vs. OCGOL-B, p=0.0003; OCGOL-A vs. OCGOL-B, p=0.1084). E) Speed of mouse during run epochs (cm/s) (one-way RM ANOVA, p<0.0001; Tukey *post-hoc* test: RF vs. OCGOL-A, p=0.0003; RF vs. OCGOL-B, p=0.0004; OCGOL-A vs OCGOL-B, p=0.9948). F) Number of laps run per session type (Friedman test, p=.0520; Dunn’s multiple comparisons *post-hoc* test: RF vs. OCGOL-A, p=0.6072; RF vs OCGOL-B, p=0.9804; OCGOL-A vs. OCGOL-B, p=0.0723). All data presented in figure: n=13 mice unless otherwise specified, means plotted with SEM.

### Task demand alters calcium event and place cell dynamics

We hypothesized that both Ca2+ transients and place cell tuning would reflect differences in task complexity and demand between the simpler RF and the multisensory OCGOL paradigms.

Throughout the animal’s behavioral progression, from RF to OCGOL, we imaged Ca2+ activity in CA1 pyramidal neurons of expressing GCaMP6f (using a Thy1-GCaMP6f transgenic mouse with uniform expression in hippocampal CA1) (Fig. 2A, 2B). Mean Ca2+ event rate and duration during run epochs were not significantly different between RF and OCGOL (Fig 2D, 2G). However, the area under the curve (AUC) and amplitude of the Ca2+ events were both increased in OCGOL compared to RF (Fig. 2E, 2F). We parsed our data based on the shape of the Ca2+ transients and GCaMP6f off-kinetics (Fig. S1A-S1D). Given the dynamics of GCaMP6f, this increase in amplitude and AUC could be due to greater burst firing of soma during OCGOL^55^.

**Figure 2:**
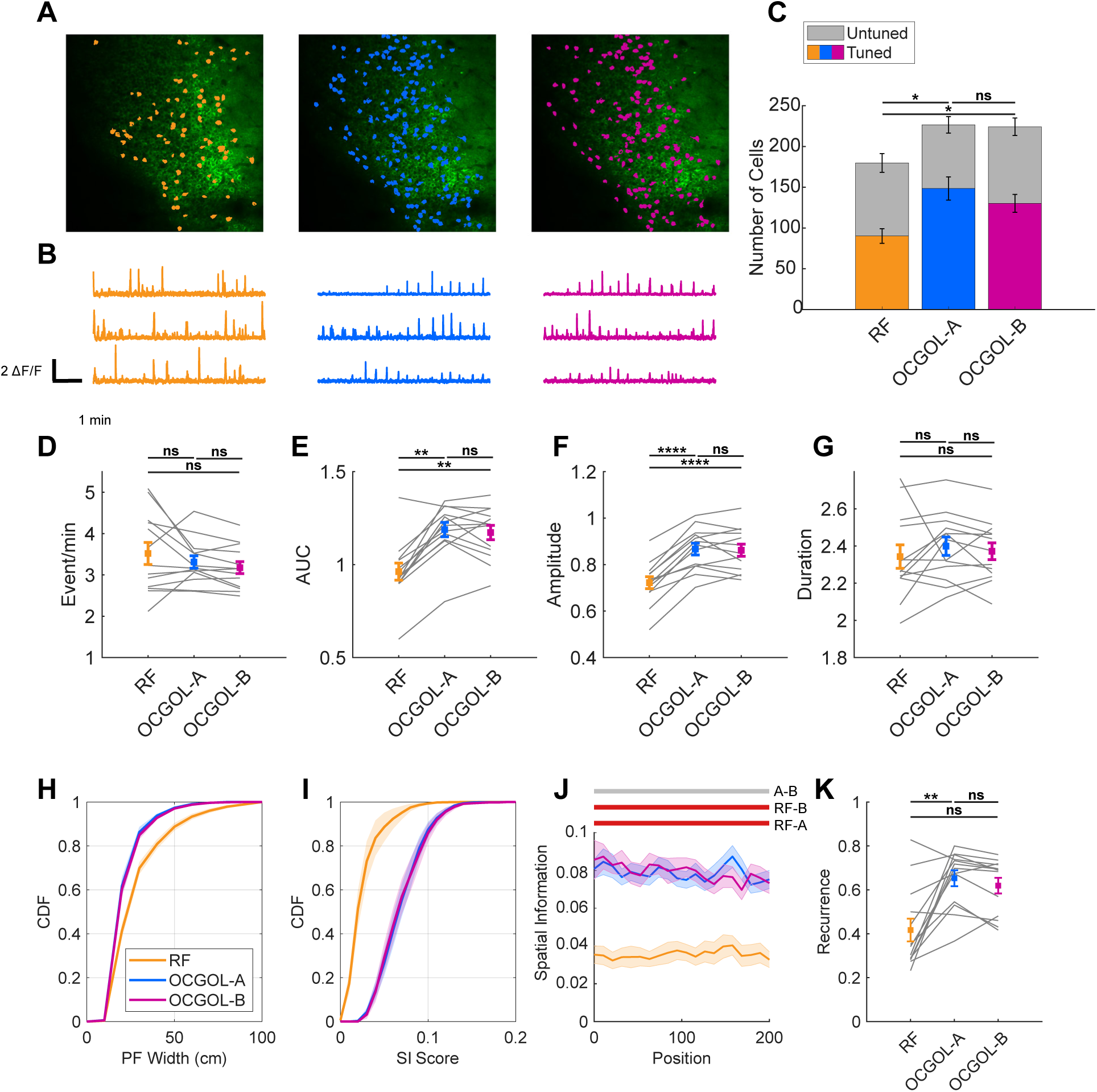
Task demand alters calcium event and place cell dynamics. A) Example field of view (FOV) of the CA1 pyramidal layer showing the mean intensity projection in a Thy1-GCaMP6f mouse during (1) random foraging (left, orange) and, 1-2 weeks later, during (2) OCGOL-A laps (middle, blue) and OCGOL-B laps (right, magenta). Overlay of neurons that met place cell criteria based on spatial information (SI). B) Example ΔF/F traces of neurons during the respective behavioral task. C) Total counts of active, untuned cells (gray) and tuned, place cells (color) averaged across all mice for each task (Total active cells: one way RM ANOVA, p=0.0109; Tukey *post-hoc* test: RF vs. OCGOL-A, p=0.0230; RF vs. OCGOL-B, p=0.0346; OCGOL-A vs. OCGOL-B, p=0.5159. Fraction of tuned cells: Friedman test, p=0.0498; Dunn’s multiple comparisons *post-hoc* test: RF vs. OCGOL-A, p=0.0558; RF vs. OGOL-B, p>0.9999; OCGOL-A vs. OCGOL-B, p=0.2327). D) Calcium event rate (events/minute) during run epochs across all tasks. Each value corresponds to the average event rate across all detected neurons for each mouse averaged across all mice (one way RM ANOVA, p= 0.1624; Tukey *post-hoc* test: RF vs OCGOL-A, p= 0.6024; RF vs OCGOL-B, p= 0.1957; OCGOL-A vs. OCGOL-B, p= 0.0596). E) AUC/event during run epochs across all tasks. Each value corresponds to the average AUC/event across all detected neurons for each mouse averaged across all mice. (Friedman test, p=0.0009; Dunn’s multiple comparisons *post-hoc* test: RF vs. OCGOL-A, p=0.0026; RF vs. OCGOL-B, p=0.0051; OCGOL-A vs. OCGOL-B, p>0.9999). F) Event amplitude during run epochs across all tasks. Each value corresponds to the average event amplitude across all detected neurons for each mouse averaged across all mice (one way RM ANOVA, p= <0.0001; Tukey *post-hoc* test: RF vs OCGOL-A, p<0.0001; RF vs OCGOL-B, p<0.0001; OCGOL-A vs. OCGOL-B, p= 0.9206). G) Calcium event duration (seconds) during run epochs across all tasks. Each value corresponds to the average duration across all detected neurons for each mouse averaged across all mice (Friedman test, p=0.0498; Dunn’s multiple comparisons *post-hoc* test: RF vs. OCGOL-A, p=0.0558; RF vs. OCGOL-B, p=0.2327; OCGOL-A vs. OCGOL-B, p>0.9999). H) Cumulative frequency of place field width (centimeters) for place cells averaged across all mice (one way RM ANOVA, p<0.0001; Tukey *post-hoc* test: RF vs. OCGOL-A, 39±1.2 vs. 31±0.85, p=0.0006; RF vs. OCGOL-B, 39±1.2 vs. 32±0.68, p=0.0008; OCGOL-A vs. OCGOL-B, 31±0.85 vs. 32±0.68, p=0.8356). I) Cumulative frequency of SI score (bits/Ca2+ event) for place cells averaged across all mice (Friedman test, p=0.0002; Dunn’s multiple comparisons *post-hoc* test: RF vs. OCGOL-A, 0.035±0.0045 vs. 0.079±0.0044, p=0.0012; RF vs. OCGOL-B, 0.035±0.00445 vs. 0.079±0.0051, p=0.0012; OCGOL-A vs. OCGOL-B, 0.079±0.0044 vs. 0.079±0.0051, p>0.9999). J) SI score per 10-cm spatial bin of the belt. Each value corresponds to the average SI across all place cells for each mouse averaged across all mice. Horizontal bars represent p-values for all pairwise task combinations for a given spatial bin by paired Wilcoxon signed rank test (gray: p>0.05, red: p<=0.05). See summary table for significant bin-wise p-values. K) Recurrence, the probability that a spatially tuned place cell maintains its identity as a place cell the subsequent day, averaged across all mice (Friedman test, p=0.0058; Dunn’s multiple comparisons *post-hoc* test: RF vs. OCGOL-A, p=0.0051; RF vs. OCGOL-B, p=0.9804; OCGOL-A vs. OCGOL-B, p=0.0930). All data presented in figure: n=13 mice unless otherwise specified, means plotted with SEM.

We found that OCGOL recruited a larger number of active CA1 pyramidal neurons compared to RF (Fig. 2G). Although there was a trend toward an increased proportion of place cells, as defined by standard spatial information (SI) score criteria, in OCGOL, we found no statistically significant differences in place cell fractions across the tasks. This suggests that task complexity increased recruitment of active cells but not specifically place cells. Similar to our previous findings^22^, place cells that were tuned to only one trial type had place fields shifted earlier on the belt, reflecting greater representation of task relevant zones (Fig. S1E-S1H). To investigate whether the overall quality of place cells was impacted by task complexity, we compared place field width and SI score of place cells between RF and OCGOL (Fig. 2H-J). In OCGOL, place field width was lower, and SI score was higher than that of RF. SI scores were higher across the belt and were not restricted to cue or reward-related zones (Fig. 2J). Further in OCGOL, place cells were more likely to be recurrent and remain a place cell across days, as opposed to losing their spatial tuning properties (Fig. 2K). These findings indicate that, while the proportion of place cells is not different between low and high complexity tasks, place cells in the higher complexity task are of higher quality and precision and are distributed in a task-selective manner. We examine this later with population vector decoding (Fig. 6F-6I).

### Task demand increases place cell and CA1 ensemble stability

Next, we investigated whether CA1 neurons and their ensemble representations were more stable in OCGOL than in RF. Our previous study established that OCGOL maintains high levels of stability across many days, but here were compare OCGOL directly to RF^22^. To assess the stability of individual cells’ tuning, we visually matched active cells between neighboring days during RF and subsequently during OCGOL. We computed the spatial tuning curves of place cells and calculated the tuning curve (TC) correlation across days for each behavioral task (Fig. 3A, 3C). Place cells in OCGOL, regardless of the trial type used for analysis, were more stable in their spatial tuning than place cells in RF. We also calculated the shift of place field centroids over time, as an alternative measure of place cell stability and, in corroboration, found reduced shift in OCGOL (Fig. 3B). Thus, place cells are better at maintaining their spatial code as task complexity increases in OCGOL.

**Figure 3:**
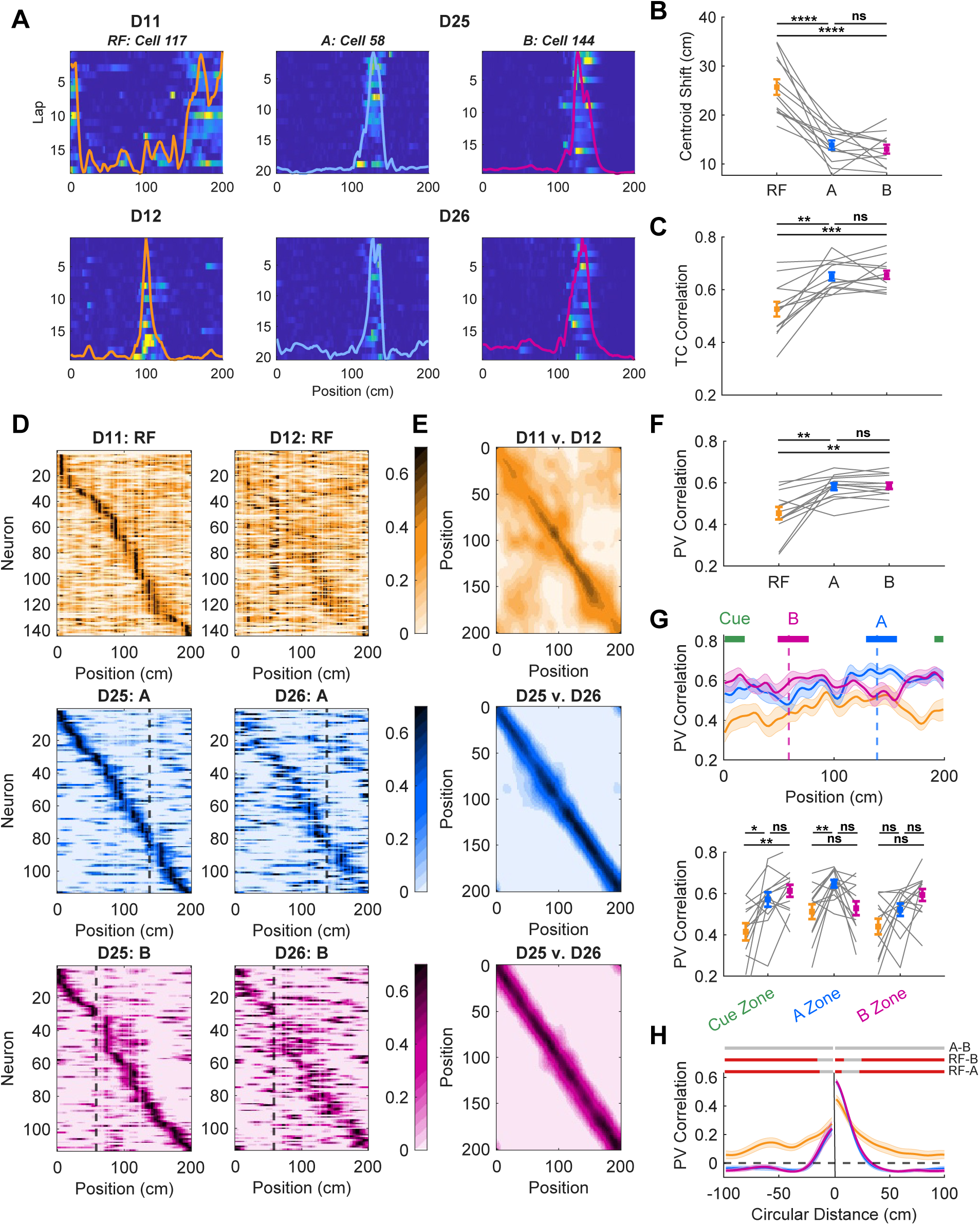
Task demand increases place cell and CA1 ensemble stability. A) Example ROIs ΔF/F plotted by lap and position for neighboring days 1 (top) and subsequent day 2 (bottom). Spatial tuning curve overlaid for each ROI in each task. B) Mean centroid shift of place cells (cm) for neighboring days (one way RM ANOVA, p<0.0001; Tukey *post-hoc* test: RF vs. OCGOL-A, p<0.0001; RF vs. OCGOL-B, p<0.0001; OCGOL-A vs. OCGOL-B, p=0.7783). C) Mean tuning curve (TC) correlation of place cells for neighboring days (one way RM ANOVA, p<0.0001; Tukey *post-hoc* test: RF vs. OCGOL-A, p=0.0023; RF vs. OCGOL-B, p=0.0002; OCGOL-A vs. OCGOL-B, p=0.9289). D) Example rate maps from a single mouse for all cells on day 1 (left) and day 2 (right) for each task. Each neuron’s rate is normalized to its maximum rate and sorted according to the position of its maximum rate across 100 spatial bins. E) Mean population vector (PV) correlation maps across all mice for each task. F) Mean PV correlation of all cells for neighboring days (one way RM ANOVA, p=0.0015; Tukey *post-hoc* test: RF vs. OCGOL-A, p=0.0060; RF vs. OCGOL-B, p=0.0039; OCGOL-A vs. OCGOL-B, p=0.8432). G) PV correlation plotted by spatial bin for each task (top). See Summary table for significant bin-wise p-values. Mean PV correlations for 30 cm zones centered at salient sensory locations (bottom): odor presentation (Cue Zone, green), reward zone B (B Zone, magenta), and reward zone A (A Zone, blue) (Cue Zone: one way RM ANOVA, p=0.0013; Tukey *post-hoc* test and paired t-test: RF vs. OCGOL-A, p=0.0289, p=0.0115 (respectively); RF vs OCGOL-B, p=0.0034, p=0.0013; OCGOL-A vs. OCGOL-B, p=0.4116, p=0.2115. Zone A: Friedman test, p=0.0079; Dunn’s multiple comparisons *post-hoc* test and paired Wilcoxon signed rank test: RF vs. OCGOL-A, p=0.0098, p=0.0081 (respectively); RF vs OCGOL-B, p=>0.9999, p=0.8926; OCGOL-A vs. OCGOL-B, p=0.0558, p=0.0110 (paired t-test). Zone B: Friedman test, p=0.1462; Dunn’s multiple comparisons *post-hoc* test and paired Wilcoxon signed rank test: RF vs. OCGOL-A, p=0.9804, p=0.1677 (respectively); RF vs OCGOL-B, p=0.1496, p=0.0143 (paired t-test); OCGOL-A vs. OCGOL-B, p=0.9804, p=0.1099). H) PV correlation of opposing spatial bins using off-diagonal values for each task (Friedman test, Effect of task p=0.00025; paired Wilcoxon signed-rank *post-hoc* test with FDR correction: RF vs. OCGOL-A: p=0.0007, RF vs. OCGOL-B: 0.0007, OCGOL-A vs. OCGOL-B: 0.89). Horizontal bars represent p-values for all pairwise task combinations for a given spatial bin by paired Wilcoxon signed rank test (gray: p>0.05, red: p<=0.05). See Summary table for significant bin-wise p-values. All data presented in figure: n=13 mice unless otherwise specified, means plotted with SEM.

To assess CA1 ensemble stability, agnostic to place cell criteria, we computed population vector (PV) correlations (Fig. 3D). We used the activity of all active cells, regardless of spatial tuning, for each spatial bin correlated against all possible spatial bins the subsequent day to create a PV correlation map (Fig. 3E). CA1 ensembles were more stable when comparing identical spatial bins across days for OCGOL compared to those in RF (Fig. 3F). We also computed PV correlation using only place cells and observed a similar increase in stability with OCGOL (Fig. S2A-S2C). We observed increased stability in salient, task-relevant zones: the odor cue zone was more stable in OCGOL than RF, and A and B-related reward zones were more stable than RF in their corresponding trial types (Fig. 3G). This relationship was also seen when restricted to place cells (Fig. S2D). Given the episodic structure and task demand of OCGOL, we also expected the converse, that comparing opposite spatial bins across days would result in lower correlations compared to that of RF. We found that when comparing PV correlations between spatial bins of increasing distance (off diagonals of the correlation map), OCGOL ensembles had anti-correlated, negative values that were significantly lower than those of RF (Fig. 3H). This relationship did not persist when restricting analysis to place cells (Fig. S2E). To confirm that this overall increased stability was specific to the OCGOL task and not due to the general experience of learning, we re-exposed a subset of mice to RF after learning and found that PV and TC scores were comparable to those of the initial RF (Fig. S4C-S4E). Overall, OCGOL increases CA1 ensemble stability and promotes discrimination between opposing spatial bins.

### Excitatory activity from lateral entorhinal cortex supports behavioral performance and task selective place cell dynamics during OCGOL

Next, we asked what could be a candidate circuit mechanism that contributes to behavioral task-driven stabilization of hippocampal place maps. Given the multimodal and odor context-based structure of the OCGOL task, we predicted that lateral entorhinal cortex (LEC), an upstream cortical region involved in contextually salient (rewards, punishments, task rules) and olfactory information processing, may support the behavioral and neural dynamics observed in this task^35,42,48^. Support for this idea comes from recent studies implicating LEC in context-dependent associative and episodic memory and driving experience dependent stabilization of place maps in area CA3^46^. To evaluate the role of LEC in this paradigm, we chemogenetically silenced LEC excitatory neurons, bilaterally, using an inhibitory DREADD (Fig. 4A, S3A). Expert mice that underwent the previous RF and OCGOL training were injected intraperitoneally for 4 days with either saline or CNO before performing randomized A and B trials in OCGOL (Fig. 4B). This paradigm allowed for tracking CA1 stability between days and conditions within each animal. When comparing behavioral performance of mice between day 2 (saline) and day 3 (CNO), we observed a slight, but significant impairment in fraction of licks within the reward zone and a consistent decrease in the fraction of correct trials in both trial types (Fig. 4C-4F). Incorrect trials were defined as trials in which the mice missed reward collection in the appropriate zone or licked in the off-target reward zone. Mouse speed upon entrance and exit of the reward zones with LEC silencing was not statistically significant from the saline injected session (Fig. S3B). Although mild, these impairments suggest that LEC plays a role in supporting recall performance of this complex context-dependent behavioral task, even at this well consolidated stage of memory.

**Figure 4:**
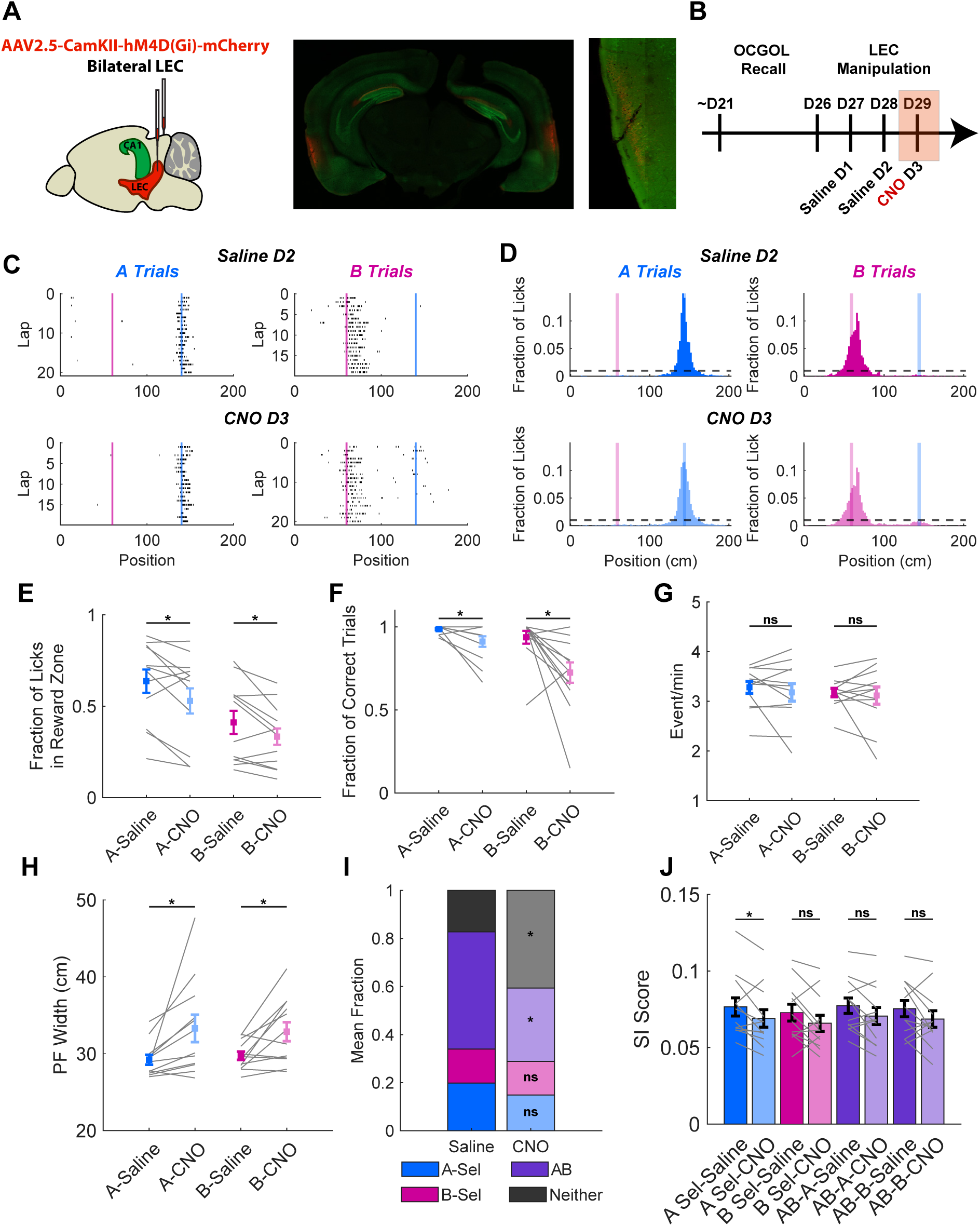
Excitatory activity from lateral entorhinal cortex supports behavioral performance and task selective place cell dynamics during OCGOL A) Schematic of CaMKII-driven inhibitory hM4D(Gi) receptor injected bilaterally into LEC (left). Sample widefield image of Thy1-GCaMP6f animal with injection site in LEC of the mCherry (red) tagged receptor (middle). Sample confocal image of LEC injection site (right). B) Schematic of behavioral timeline for silencing at recall of OCGOL. Shaded red rectangle denotes the day of LEC silencing with i.p. CNO. C) Example lap by lap lick raster plot for a single mouse with saline (top row) and with CNO injection (bottom row). Magenta vertical line denotes the start of the B reward zone and blue denotes the start of the A reward zone. D) Mean fraction of licks per 2-cm spatial bin of the treadmill belt across all mice in A and B trials with saline (top row, blue and magenta) or CNO (bottom row, light blue and light magenta). E) Fraction of licks in A or B reward zones with saline (blue, magenta) or CNO (light blue, light magenta) (Paired t-test, A-Saline vs. CNO, p= 0.0300; B-Saline vs. CNO, p=0.0306). F) Fraction of correct trials in A or B trials with saline or CNO (paired Wilcoxon signed rank test, A-Saline vs. CNO, p= 0.0312; B-Saline vs. CNO, p=0.0112). G) Calcium event rate (events/minute) during run epochs with saline or CNO (Paired t-test, A-Saline vs. CNO, p= 0.4056; B-Saline vs. CNO, p=0.6878). H) Place field width (cm) for place cells with saline or CNO (Paired t-test, A-Saline vs. CNO, p= 0.0218; B-Saline vs. CNO, p=0.0127). I) Fraction of cells that are tuned in A (blue), B (magenta), A and B (purple), or not tuned in either (black) with saline or CNO (light blue, light magenta, light purple, gray) (Paired t-test, A-selective Saline vs. CNO, p= 0.1176; B-Selective Saline vs. CNO, p= 0.9277; AB Shared Saline vs. CNO, p= 0.0195; paired Wilcoxon signed rank test, Untuned Saline vs. CNO, p=0.0161). J) SI score (bits/Ca2+ event) for cell types with saline or CNO (Paired t-test, A-Saline vs. CNO, p= 0.0346; B-Saline vs. CNO, p=0.1078; AB Shared in A Trials Saline vs. CNO, p=0.0537; AB Shared in B Trials Saline vs. CNO, p=0.1432). All data presented in figure: n=12 mice unless otherwise specified, means plotted with SEM.

Next, we examined how LEC silencing affects the task relevant neural dynamics and spatially tuned activity. We found that silencing of excitatory activity in LEC did not significantly alter the number of active cells (Fig. S3C) or the rate of calcium events during run epochs in active cells (Fig. 4G). Restricting analysis to only significantly tuned place cells, we found a drop in the overall fraction of tuned cells with their place field width significantly increased upon silencing of LEC (Fig. 4H, 4I). In our previous work, we identified trial-specific task-selective neurons that emerge during OCGOL learning and recall^22^. Therefore, we classified cells as A-selective (only tuned in A), B-selective (only tuned in B), AB shared (tuned in both trial types), and neither (not tuned in either trial type). With LEC silencing, the proportion of AB shared cells significantly decreased, and correspondingly, the proportion of neither (untuned) cells significantly increased (Fig. 4I). Thus, the overall number of place cells is reduced with LEC silencing, specifically due to a reduction in the place cell population tuned to both contexts. Next, we compared the quality of place cells, given their observed higher SI scores in the OCGOL. We found that SI scores decreased in A trials for A-selective place cells (p <0.05) and AB shared place cells (p=0.0537) (Fig. 4J). Together, these observations suggest that reducing LEC excitatory drive decreases the proportion of significantly tuned place cells, and the remaining place cells are broader in their tuning fields. The trial specific decrease in SI score may be due to the higher demand of navigating to the far reward zone.

### LEC silencing disrupts neural and ensemble stability in OCGOL

To assess whether LEC excitatory populations support the high levels of stability seen in OCGOL, we compared TC and PV correlation between neighboring days without LEC silencing, day 1 (saline) vs. day 2 (saline), and with LEC silencing, day 2 (saline) vs. day 3 (CNO). TC correlation between days was significantly reduced with LEC silencing (Fig 5A, 5C). Likewise, the degree of place field centroid shifts increased significantly with LEC silencing compared to without, but this only reached statistical significance for A trials (Fig. 5B). We plotted behavioral performance against TC correlation for each mouse and found that, prior to LEC silencing, both measures are uniformly high across mice at recall stage; suggesting a ceiling effect (Fig. 5D). However, with LEC silencing, we observe TC correlation and behavioral performance become positively correlated with one another across a dynamic range (Fig. 5E). This spread may be representative of the animal-to-animal variability in degree of LEC suppression. This reached statistical significance for B trials, but not for A trials, which is expected as B performance was preferentially affected compared to A. This demonstrates that LEC activity supports CA1 spatial coding and that, even at memory recall, CA1 activity is still correlated with behavioral performance. PV correlation scores taken across matched cells between days significantly decreased with LEC silencing (Fig. 5F, 5G). We found that PV scores were similarly correlated to behavioral performance (Fig. S4A, S4B). When PV correlations were parsed by position on the belt, there was no clear location that was most susceptible to LEC perturbation (Fig. 5H). Much of the track decreased in ensemble stability after silencing, indicating the global impact of LEC across space. Further, CA1’s dependence on LEC for stability seems to be specific to OCGOL, as no significant deficit was observed when we silenced LEC in RF for a subset of mice (Fig. S4C-S4E). Finally, we examined the PV correlation between A and B laps within the same day for the AB shared population of cells, the population we earlier found to be most vulnerable to LEC silencing, as opposed to one trial type across days (Fig 5I-5K). Typically, in OCGOL, the spatial representations between A and B laps are orthogonal, with low PV correlation scores between trial types^22^. They are especially low at the sites of the reward zones—this is conceivably where the track is most behaviorally different for the mouse. With LEC silencing, we observed that PV correlation score between A and B trials significantly increased, meaning that the spatial representations between the two contrasting trial types became more similar (Fig. 5J). When separated by position, LEC silencing significantly impacted the first third of the belt (Fig. 5K). This area contains the odor cue and the track cues prior to the B reward zone. This suggests that there is a greater neural overlap at the point in space where the animal needs to decide its trial-specific performance. Taken together, silencing of LEC excitatory cells reduces overall place cell and ensemble level stability across days as well as discrimination between contexts within days.

**Figure 5:**
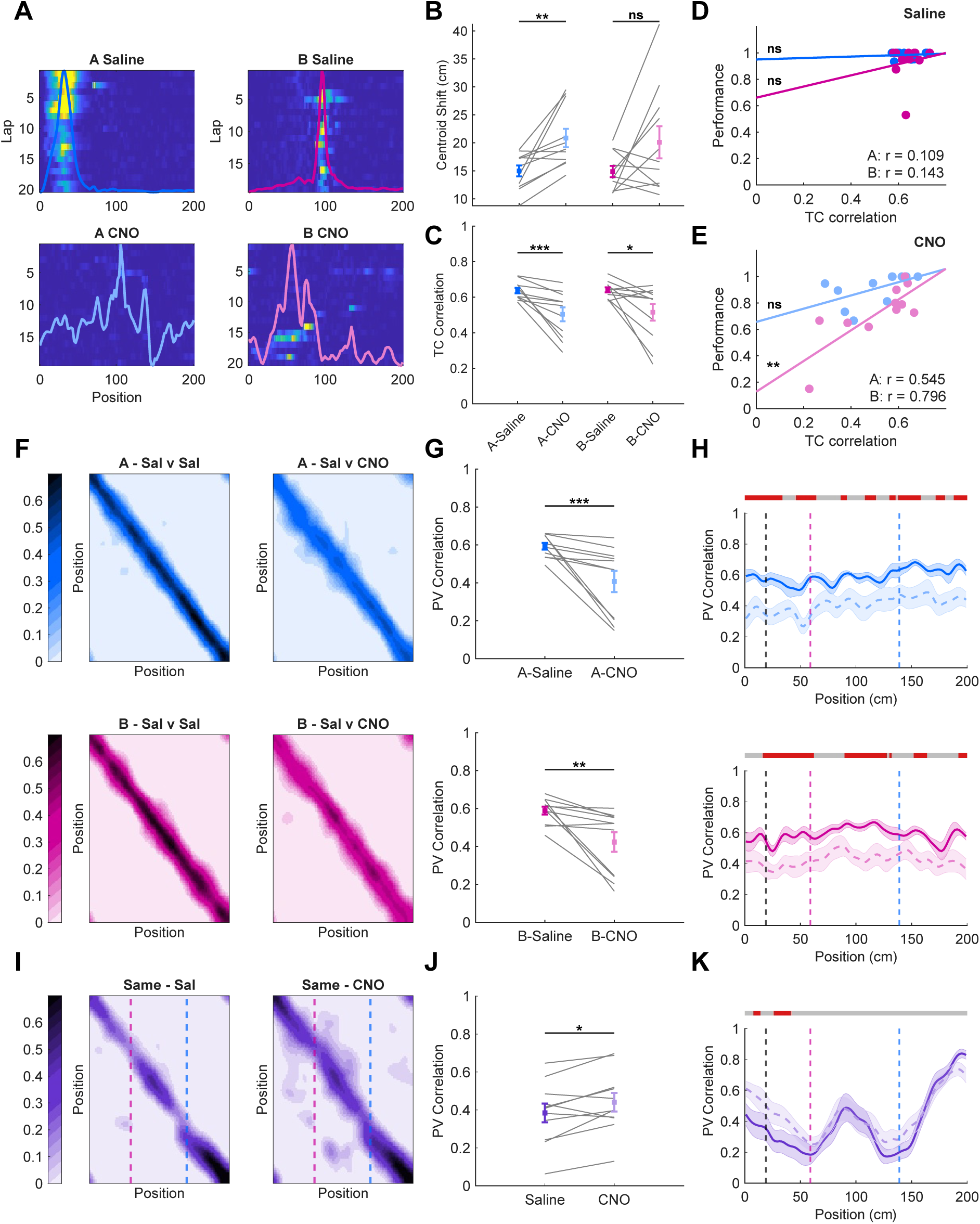
LEC silencing disrupts neural and ensemble stability in OCGOL. A) Example ROIs ΔF/F plotted by lap and position with saline (top) or CNO (bottom). Spatial tuning curve overlaid for each ROI in A trials (left, blue/light blue) and B trials (right, magenta/light magenta). B) Mean centroid shift of place cells (cm) for neighboring days: day 2 saline and day 3 CNO (n=10, A-Saline vs. CNO: Paired t-test, p=0.0053; B-Saline vs. CNO: Paired t-test, p=0.0943). C) Mean TC correlation of place cells with saline or CNO (paired Wilcoxon signed rank test, A-Saline vs. CNO, p=0.0010; B-Saline vs. CNO, p=0.0186). D) TC correlation (day 1 saline vs. day 2 saline) plotted against behavioral performance (day 2 saline) during A (blue) and B trials (magenta) (Two-sided Pearson correlation, A: r=0.109, p=0.749; B: r=0.143, p=0.675). E) TC correlation (day 2 saline vs. day 3 CNO) is plotted against behavioral performance (day 3 CNO) (Two-sided Pearson correlation, A: r=0.545, p=0.083; B: r=0.796, p=0.003). F) Mean PV correlation map between day 1 saline and day 2 saline (left column) and between day 2 saline and day 3 CNO (right column) for A trial (blue) and B trials (magenta) across all mice. G) Mean PV correlation of all cells with saline or CNO (paired Wilcoxon signed rank test, A-Saline vs. CNO, p=0.0010; B-Saline vs. CNO, p=0.0029). H) PV correlation plotted by spatial bin with saline (solid) or CNO (dashed) for A trials (top) and B trials (bottom), Cue and reward zones denoted by vertical lines. Horizontal bars represent p-values for a given spatial bin by paired Wilcoxon signed rank test (gray: p>0.05, red: p<=0.05). See Summary table for significant bin-wise p-values. I) Mean same day PV correlation map, for cells tuned in both trial types, between A and B trials with saline (left) or CNO (right) across all mice. J) Mean same day PV correlation between A and B trials with saline (purple) or CNO (light purple) for AB shared cells (Paired t-test, p=0.0261). K) Same day PV correlation plotted by spatial bin with saline (solid) or CNO (dashed). Horizontal bars represent p-values for a given spatial bin by paired Wilcoxon signed rank test (gray: p>0.05, red: p<=0.05). See Summary table for significant bin-wise p-values. All data presented in figure: n=11 mice unless otherwise specified, means plotted with SEM.

### LEC silencing and reduced task demand decrease performance of a decoder

To further probe context discrimination with LEC silencing, we constructed a population vector decoder, trained on activity from the first half of the session, to predict both context/trial type (A or B lap) and position of the animal within a given session (Fig. 6A-6B). With LEC silencing, the decoder had significantly more errors in predicting the current trial (identification error) (Fig. 6D). Notably there was a significant change in number of errors at the reward zones, sites where there are typically lower likelihoods of error (Fig. 6C). In addition to context, LEC silencing also significantly increased errors in position decoding (Fig. 6E, 6I).

**Figure 6:**
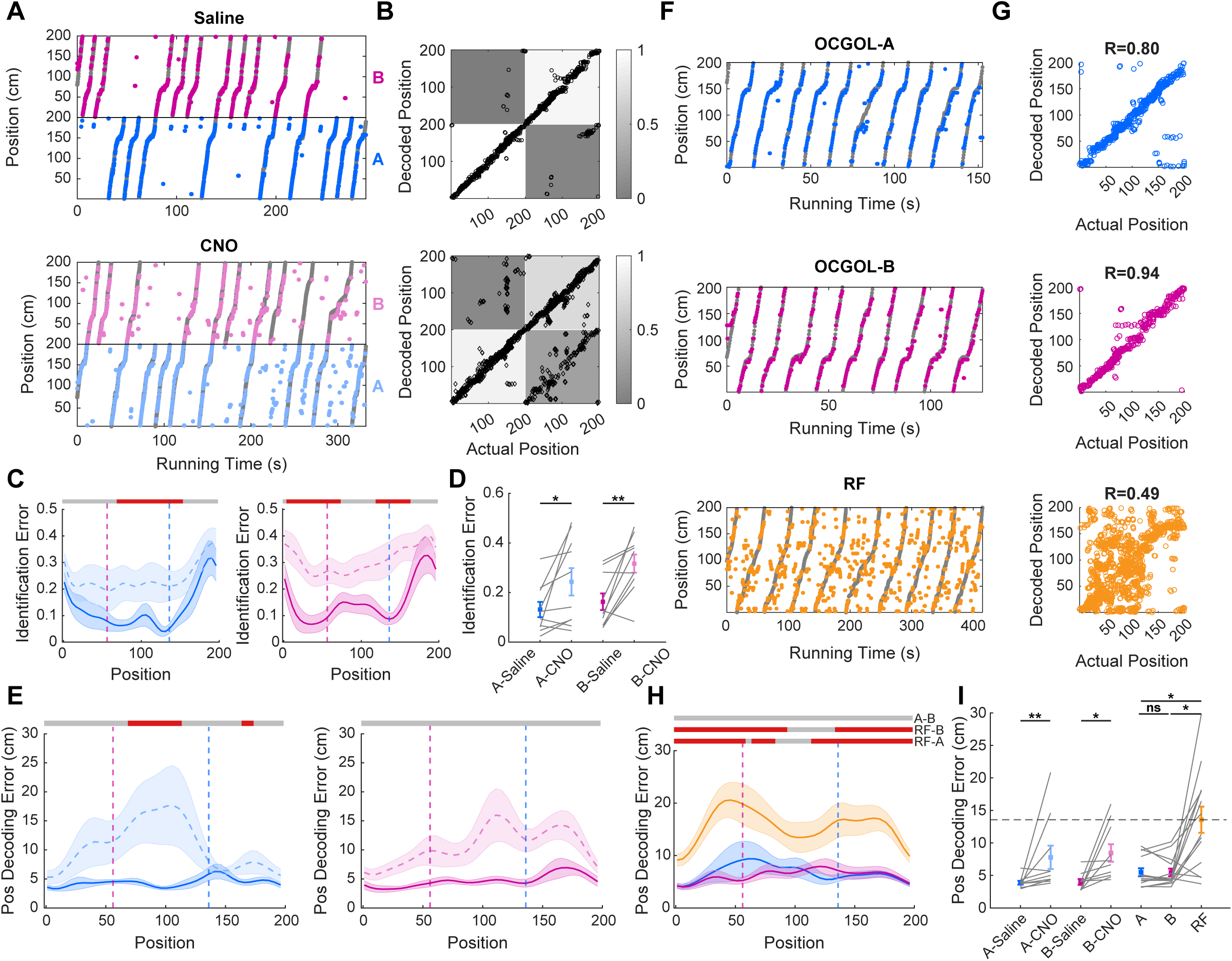
LEC silencing and reduced task demand decrease performance of a decoder. A) Visualization of a PV decoder’s predictions with saline (top) or CNO (bottom). Decoded positions (A trial-blue, B trial-magenta) are plotted on top of actual positions (gray) as a function of time. Position 0-200 appear twice, once for each trial type. B) Sample confusion matrix of a PV decoder predicting both an animal’s position and trial type with saline (top) or CNO (bottom). Black points are plotted according to the animal’s actual position and trial against the decoder’s predictions. Each cell represents the number of decoded points within that quadrant divided by the total data points for each trial type. C) Trial identification error plotted by spatial bin with saline (solid) or CNO (dashed) for A and B trials. Reward zones are plotted in vertical dashed lines. Horizontal bars represent p-values for a given spatial bin by paired Wilcoxon signed rank test (gray: p>0.05, red: p<=0.05). See Summary table for significant bin-wise p-values. D) Mean trial identification error with saline or CNO for A and B trials (Paired t-test, A-Saline vs. CNO, p=0.0428; B Saline vs. CNO, p=0.0048). E) Position decoding error plotted by spatial bin with saline (solid) or CNO (dashed) for A and B trials. Horizontal bars represent p-values for a given spatial bin by paired Wilcoxon signed rank test (gray: p>0.05, red: p<=0.05). See Summary table for significant bin-wise p-values. F) Example position decoding for a mouse across OCGOL-A (top, blue), OCGOL-B (middle, magenta), and RF (orange). G) Sample scatter plot of actual position against decoded position for an animal across all tasks. H) Position decoding error plotted by spatial bin for RF and OCGOL. Horizontal bars represent p-values for all pairwise task combinations for a given spatial bin by paired Wilcoxon signed rank test (gray: p>0.05, red: p<=0.05) (n=14). See Summary table for significant bin-wise p-values. I) Position decoding error for animals that underwent LEC silencing (left four columns) and those that completed RF and OCGOL (right three columns). Dashed horizontal gray line at the mean position decoding error during RF (A-Saline vs. CNO: Wilcoxon signed rank test, p=0.0059; B-Saline vs. CNO: Paired t-test, p=0.0168; RF vs. OCGOL comparison: n=14, Friedman test, p=0.0128; Dunn’s multiple comparisons *post-hoc* test: RF vs. OCGOL-A, p=0.0421; RF vs OCGOL-B, p=0.0245; OCGOL-A vs. OCGOL-B, p>0.999). All data presented in figure: n=10 mice unless otherwise specified, means plotted with SEM.

To more directly implicate the role of LEC in specifically supporting neural dynamics underlying higher complexity behavioral tasks (OCGOL), we postulated that the lower demand, simpler RF task is unlikely to engage LEC. Accordingly, we predicted that the low complexity task, RF, would have similar levels of decoding errors as those with LEC silencing. Since there were no alternative contexts within the RF task, we only performed population vector decoding for position (Fig. 6F-6G). We found that RF had significantly higher decoding error compared to equivalent laps run in OCGOL (Fig. 6H-6I). Given our earlier finding that OCGOL recruits a greater number of active neurons, and to ensure fair comparison, we trained the OCGOL decoder on a random subsample of ROIs to match the number of ROIs used for RF per mouse (Fig. S5A-S5D). We found that the increase in decoding error for RF persisted even with the difference in the number of ROIs accounted for – likely due to the decreased place cell SI score we observed earlier. However, the position decoding error during RF was still significantly greater than that seen with LEC silencing (Fig. 6I). This intermediate level of decoding error with LEC silencing was not too surprising as the perturbation occurs during recall, the task is likely supported by other brain regions in conjunction with LEC, and because LEC activity is suppressed, but not completely removed with the viral approach. Nevertheless, our observed effects of LEC silencing on task performance, place map stability, and spatial and contextual decoding in OCGOL behavior implicate LEC in supporting task-relevant neural dynamics during recall phases of memory-guided behavior. Overall, with the findings that place cell numbers, quality, and stability increase with task complexity, we propose that LEC is likely engaged by higher complexity associative learning behaviors and contribute to behaviorally driven context-dependent expansion and maintenance of place maps.

## Discussion

In this study, we probed behavioral and neural circuit mechanisms to address the multifaceted stability-plasticity literature^15,16,21,24,56^. While neural representations must remain dynamic to support adaptive learned behaviors, stabilization of neural assemblies is critical for memory retention. To resolve this conundrum, we postulated that CA1 ensembles stabilize as a function of cognitive demand and that LEC facilitates this stability resulting in reliable memory recall. Our study design allowed us to interrogate the effect of varying cognitive demand on CA1 neural dynamics, within the same animals, as they navigated in a fixed physical environment. The low demand task had no predictable reward zones, and animals had to randomly forage to seek rewards, whereas the high demand task had fixed, context-dependent reward zones that the animals learned to navigate to (Fig. 1A-1D). We found that a greater number of cells were recruited in the high demand task and with larger Ca^2+^ transients (Fig. 2C, 2E, 2F). Of those cells active in the high demand task, the spatially tuned place cells had greater information content and more precise tuning than those in the low demand task (Fig. 2H-2J). Individual place cells and whole CA1 ensemble representations were more stable between days in the high demand task than in the low random foraging task (Fig. 3C, 3F). During OCGOL, the stability of task-relevant zones was preferentially boosted (Fig. 3G). These findings systematically confirm that sensory-spatial context driven behavioral demand can enhance CA1 place coding and bolster its stability between days. Given the stability we observed in the high demand, odor-contextual navigational task, we tested whether upstream LEC, an area known to encode odor, novelty, and learning rules, contributes to this stability through a loss of function chemogenetic silencing approach. We found that silencing LEC during long term recall of the high demand task not only impaired mouse performance, but reduced place cell proportion, and increased place field width (Fig. 4F, 4H, 4I). Place cells tuned in both A and B trials were most affected. Their overall proportion was reduced and their discrimination between contexts was impaired (Fig. 4I, 5J, 5K). Moreover, there were even greater effects seen on day-to-day stability with a significant reduction in both PV and TC correlations which were linearly correlated with performance (Fig. 5C, 5E, 5G). Our findings demonstrate that a high-demand odor-contextual task helps stabilize an otherwise dynamic hippocampal representation of space and that this stability relies on input from LEC even during memory recall.

### Task demand and place coding accuracy

With the introduction of the odor-context task contingencies for the high demand task, we observed an overall refinement in CA1 coding when directly compared to coding during the low demand task—narrowed place field widths and increased SI scores. Given this increase in quality, when we trained a PV decoder on the same number of neurons in the low and high demand task, decoding performance was higher for the high demand task (Fig. S5D). These findings together validate the idea that high quality spatial maps emerge over the course of learning, organize themselves based on task-relevant cues, and remain stable during memory recall.

### Behaviorally driven representational stability

Our results further support the hypothesis that attentional demand stabilizes hippocampal ensembles to support memory recall^24^. We observed high levels of stability at both the place cell and ensemble level. Further, task-relevant spatial bins exhibited greater stability in the high demand task and ensemble maps for each spatial bin were more distinct from one another. We compared this directly in our study with random foraging which we observed to have significantly lower stability. This lower stability has been observed independently in similarly unstructured tasks and when engagement is lower during non-operant reward delivery or internal state^7,15,16,23^.

### Cortico-hippocampal interactions in modulating ensemble stability and flexibility

Canonically, LEC has been described as the non-spatial input to CA1 and only marginally influential on place coding^32,57,58^; however, growing work suggests that as tasks become more demanding and complex, LEC, in addition to the more classically implicated MEC, may actively support spatial memory recall^34,52,59^. LEC has direct projections to CA1 and indirect connections to CA1 through the hippocampal circuit and interactions with MEC, making it a promising driver of CA1 coding^57,60–62^. Chemogenetically silencing LEC at recall of the contextual task mildly but significantly impaired behavioral performance. Current literature has only implicated LEC in the learning of new non-spatial and spatial associations, so we were surprised to observe any behavioral deficits at long term memory recall^38,42,46,48^. We posit that the randomized, episodic task structure, which melds both odor-based context association and spatial navigation, engages LEC even at memory recall.

Our chronic calcium imaging further confirmed CA1’s dependence on LEC with significant deterioration of stable place maps. Interestingly, it also highlighted a neural deficit in context discrimination - place cells that were active in both A and B trials, typically orthogonal in their coding, showed greater similarity between trial types. This deficiency was further reflected by diminished PV decoding of trial type with notably higher rates of error at reward zones, areas that are typically very good at context decoding. These findings bolster the role of LEC in CA1 place coding, given a complex task structure, and further implicate LEC in context-dependent remapping alongside studies conducted in other hippocampal subregions^45,46,51^. Silencing LEC in the high demand task at recall did not completely drop PV position decoding errors to RF levels, but this is unsurprising given that CA1’s activity will still be supported by other synaptic inputs and potentially consolidated to cortical areas^63–65^. While our work emphasizes that LEC supports CA1 place coding and behaviorally driven ensemble stability, other parallel information processing circuits could also contribute.

### Cellular and circuit mechanisms for context-dependent place coding and stability

Our finding that LEC silencing reduces CA1 stability fits well within the framework of LEC-centric cortico-hippocampal and neuromodulatory circuit mechanisms gating associational synaptic plasticity and context-place coding. The excitatory drive from LEC is particularly well suited to facilitate dendritic spikes and non-linear integration of different synaptic inputs by virtue of the disinhibitory GABAergic microcircuits it recruits ^46,51,66^. Dendritic spikes are a key cellular substrate supporting the induction of associational plasticity rules such as input timing dependent plasticity (ITDP) and behavioral time scale dependent plasticity (BTSP) in CA1^67,51,68^ and have been functionally implicated in feature selective tuning^69–71^, including place field generation^72,73^. *In vivo,* processing of LEC-driven activity and how it engages different hippocampal microcircuits to shape stability and plasticity dynamics of neural representations will depend on the behavioral and neuromodulatory state of the animal. One of the many neuromodulators implicated in attentional demand and novelty is dopamine^24,74–76^; this system may play a role in our high-demand task. In addition to direct targeting of the hippocampus, the DA system particularly modulates LEC, via the ventral tegmental area, to encode cue-reward associations^49,51,66^. DA-gated increased LEC drive and dendritic spikes could promote long-term plasticity^77–80^, leading to the increased hippocampal stability seen in CA1 with high task demand^24,74,81^.

In conclusion, our results show that increased task demand refines individual place cell tuning and stabilizes CA1 ensembles across days. This established population stability is reduced when excitatory cells in LEC are silenced. Thus, LEC supports behaviorally-driven performance and CA1 stability at recall of a spatial-contextual task. Our findings hold translational relevance as it is well established that LEC is affected early in Alzheimer’s Disease (AD) with later synaptic spread to downstream hippocampal areas ^82–85^. Thus, future studies exploring cellular and circuit mechanisms engaged by LEC-CA1 functional interactions under behavioral demand will drive insights into the mnemonic processes underlying neural representational stabilization and identify new therapeutic targets to alleviate memory and ensemble deficits in AD.

## Methods

### Experimental model and subject details

#### Mice

Experiments were performed with 2- to 4-month-old male (n=13) and female (n=3) mice on a C57BL/6J background transgenically expressing GCaMP6f from the Thy1 locus (GP5.5 JAX strain #024276)^86^. All experiments were approved by the Institutional Animal Care and Use Committee at New York University Medical Center.

### Method details

#### Viral injections

For silencing of lateral entorhinal cortex (LEC), AAV2.5-CaMKIIa-hM4D(Gi)-mCherry was injected bilaterally in LEC. Mice were anesthetized with inhaled isoflurane (5% induction, 1.5-3% maintenance, Matrx VIP 3000 calibrated vaporizer) and buprenorphine (0.1 mg/kg i.p.) during surgery. Mice were placed in a stereotactic apparatus (Stoelting) and an incision was made in the skin to expose the skull. The skull was leveled using bregma and lambda to align the z-axis. A small craniotomy was made to expose the brain for viral injection through a glass pipette (Drummond Scientific) at the following sites LEC: A/P 3.2 and 3 from Bregma, M/L 4.5 and 4.7 from midline, D/V 2.5, 2.8 and 3.0 from brain surface (6 sites). A total of 46 nL (23 nl intervals, 15 seconds apart) of virus was injected per site (Drummond Scientific Nanoject II). Between sites with different z-coordinates there was a 2-min pause before pipette retraction. Between sites with different xy-coordinates there was a 10-min pause to allow for appropriate dispersion of the virus before slow retraction of the glass pipette out of the brain. The alternate hemisphere was then injected. The incision was sutured (Henry Schein) and antibiotic ointment (Neosporin) was applied. Mice were injected with 0.5 of saline solution subcutaneously to rehydrate, removed from anesthesia, and monitored until ambulatory. For 3 days postoperatively, mice were given buprenorphine (0.1 mg/kg, i.p.) Infection sites were confirmed post hoc. Mice that did not display viral expression in the hemisphere ipsilateral to the cranial window or had spread into ventral CA1 were excluded from LEC silencing analysis.

#### Histology

On the last day of behavior, mice were anesthetized with inhaled isofluorane (5%) and ketamine/xylazine (i.p.) until adequately sedated with no toe pinch reflex. Mice were transcardially perfused manually, using a 30 mL syringe, with 20 mL of cold phosphate-buffered saline followed by 20 mL of cold 4% paraformaldehyde (PFA) in PBS. Brains were extracted and fixed in 4% PFA in PBS at 4°C overnight. Brains were washed in 0.3M Glycine in PBS for 20 min followed by 3, 15-min washed in PBS. Brains were cut (Leica VT1000S) into 100 μm coronal slices and then mounted onto microscope slides using Vectashield Hard Set Mounting Medium with DAPI (Vector Laboratories). Slices were stored at 4°C. Brain sections were imaged using 10X epifluorescence imaging (Olympus VS200) using EGFP/FITC (green) and mCherry/TxRed (red) filters. Images were processed using QuPath and the ABBA software package^87^.

#### Hippocampal window and headpost implantation

1 week after viral injections mice were implanted with a circular imaging window (3.0 mm × 1.5 mm [diameter × height]) centered at 2.3 mm AP and 1.5mm ML over the left (n=15) or right (n=1) dorsal-intermediate hippocampus. A ∼3-mm craniotomy was drilled, the skull fragment was removed, and a vacuum system was used to aspirate the overlying cortex until reaching the external capsule. Ice-cold ACSF was used to irrigate the area throughout the duration of the aspiration procedure. An imaging window made up of a 3-mm diameter glass coverslip (64–0720 Warner) attached to a 1.5-mm deep stainless-steel cannula with optical glue (NOA-61, Norland products) was placed into the craniotomy so that the top of the cannula was flush with the skull. The cannula was secured to the skull with Vetbond. A 3-D printed headpost was attached to skull with dental cement (C&B Metabond) for head fixation.

#### In vivo two-photon imaging acquisition

Two-photon calcium imaging experiments were conducted *in vivo* using a resonant/galvanometric laser-scanning microscope (Ultima, Bruker). The system was coupled to a tunable Ti:Sapphire pulsed laser (Mai Tai DeepSee, Spectra Physics) operating at an 80 MHz repetition rate and <70 fs pulse width with dispersion compensation. Excitation light was tuned to 920 nm to drive GCaMP fluorescence. Imaging was performed through a 16x, 0.8 NA water-immersion objective (Nikon) mounted on a movable objective nosepiece. Image sequences were acquired at approximately 30 frames per second, using a frame size of 512 x 512 pixels and 1.5x digital zoom, corresponding to a field of view measuring roughly 555 x 555 µm. Fluorescence was detected with high-sensitivity GaAsP photomultiplier tubes (7422PA-40; Hamamatsu).

#### Behavioral apparatus

Mice ran on a ∼2-m treadmill track enriched with micro-textures, tones, and odorants; behavioral data (position, lap onset, sensory cues, and licking) was acquired as previously described^22^. All cues were triggered by RFIDs attached to the treadmill belt and odorant scavenging and pressure were controlled by a vacuum system.

#### Random foraging

Following recovery from surgery (3–5 days), mice were water deprived and habituated to handling and head-fixation to behavioral apparatus. Mouse weight was monitored daily to ensure it remained at least 80% of baseline. Over 2 weeks, mice were trained to operantly collect 5% sucrose water at lap-randomized water reward zones as previously described^22^. Once mice attained a running speed greater than ∼1 lap/min, they performed random foraging (RF) for two days with each session containing ∼20 laps. These days make up the RF sessions used for comparison to odor-cued goal oriented learning.

#### Odor-cued goal oriented learning

After completion of RF, mice learned odor-cued goal oriented learning (OCGOL) according to the accelerated training regimen previously described ^22^. Briefly, over the course of 1.5-2 weeks, mice learned to navigate to a single 10-cm reward zone whose position was dependent on the odorant puffed at the mouse’s snout at a 20-cm zone at the start of each lap (Odor A-10% pentyl acetate in mineral oil, Odor B-10% +-pinene in mineral oil; Reward zone A-140 cm, Reward zone B-60 cm). Each lap was cued with a 4 kHz tone for 0.5 s prior to lap start. Mice were trained until behavioral performance was >80% when trial types were randomized for two consecutive days and ∼40 laps were run. Correct trials were defined as laps in which the mouse collected rewards in the correct reward zone and suppressed licking in the incorrect reward zone. After 2 days of high performance, mice entered recall stage and performed OCGOL with randomized trials for 5 days. Recall days 4 and 5 were used for all comparisons between RF and OCGOL.

#### LEC silencing

Following 5 days of OCGOL, mice completed 4 more days of randomized OCGOL. Each day mice received an injection of either saline or CNO (5 mg/kg, i.p.) 20 minutes prior to the imaging session. After injection they recovered in their home cage. Days 1 and 2 mice were injected with saline, day 3 with CNO to silence LEC excitatory cells, and day 4 saline was injected again as a recovery day. For all comparisons between control and LEC silencing, day 2 (saline) and day 3 (CNO) are used. For all stability measures (PV and TC correlation) control values were calculated between day 1 (saline) and day 2 (saline) and silencing values were calculated between day 2 (saline) and day 3 (CNO).

#### Random foraging post-learning

For post-recall RF experiments, mice were exposed to a novel treadmill belt with an equivalent number of textures as the initial belt. They underwent 6 days of RF with randomized water reward zones on each lap to familiarize themselves with the belt. For days 1-5 mice received an injection of saline. On day 6 mice received CNO (5 mg/kg, ip). For all stability measures (PV and TC correlation) control values were calculated between day 4 (saline) and day 5 (saline) and silencing values are calculated between day 5 (saline) and day 6 (CNO). For each session, mice were required to run ∼20 laps.

### Quantification and statistical analysis

#### Image processing and signal extraction

All imaging data was motion-corrected using the NoRMCorre non-rigid motion correction algorithm implemented in MATLAB^88^.The RF day 2 session prior to learning of OCGOL, was used as the template against which all future sessions were motion corrected. For exceptions in which the RF session was not a representative image of all future sessions, an intermediate day was used as the template. All sessions that were compared to one another were motion corrected using the same template to allow for matching of ROIs. ROI detection was done using a constrained non-negative matrix factorization (CNMF) approach through the CaImAn software package^89,90^. Non-somatic ROIS were discarded through manual screening of detected ROIs. ROIs were matched between days using the register_multisession.py function in the CaImAn package – matches were then manually screened for high-quality matches.

As previously described^22^, for each ROI, the baseline fluorescence (F₀_baseline) was estimated by computing the median (50th percentile) of the fluorescence trace within a sliding 15-s window at every time point, implemented using the prctfilt.m function in CaImAn^89^. An identical sliding-window percentile approach was applied to the background component to obtain F₀_background.

Relative fluorescence changes were then calculated as:

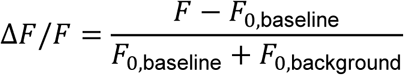

The resulting ΔF/F traces were smoothed with an exponential filter (time constant τ = 0.2 s) to attenuate photon shot noise introduced during image acquisition.

Calcium events were detected using a noise-symmetry approach in which positive and negative baseline fluctuations were assumed to occur at similar rates. Candidate events were defined as consecutive frames rising above 2 SD and ending below 0.5 SD of the mean. Events were binned by amplitude and duration, and only positive events with a false-positive rate below 5% were retained. After masking detected events, the baseline was recalculated and detection repeated twice to improve sensitivity. Events shorter than 1 s were excluded.

#### Data analysis

##### Run epochs

As described previously^7,22,25^, we defined running epochs as consecutive frames during which the mouse was moving forward with a minimum peak speed of 5 cm/s for at least 1 s in duration. Neighboring run epochs separated by less than 0.5 s were merged.

##### Calcium event dynamics

For all calcium events during run epochs, area under the curve (AUC) was calculated as the cumulative area from event onset to event offset. Event amplitude was calculated as the difference between the peak ΔF/F from ΔF/F at event onset. Event rate was calculated by dividing the total number of events during run epochs, per ROI, by the total amount of time spent running.

##### Place tuning and dynamics

###### Spatial information

Spatially tuned neurons were identified by comparing observed spatial information to a shuffle-based null distribution^7,25^. Spatial information was calculated by^91^:

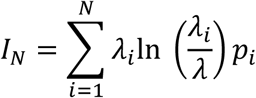

where 𝜆_𝑖_is the running-related transient rate in spatial bin 𝑖, 𝑝_𝑖_ is the occupancy probability of that bin, 𝜆 is the overall transient rate, and 𝑁 is the number of spatial bins. Transient rates were computed as the Gaussian-smoothed (𝜎= 3 bins) onset count per bin divided by occupancy time. Information was calculated across multiple spatial resolutions (𝑁= 2–100 bins). To assess significance, transient onset times were randomly reassigned within running epochs 1,000 times for each binning scheme to generate shuffled information values. To correct for binning bias, the mean shuffled information was subtracted from the observed value for each 𝑁. For each neuron, the maximum bias-corrected information across bin sizes was used as the final spatial information estimate and compared to the corresponding distribution of shuffled maxima. Cells with p < 0.05 were classified as place cells.

###### Spatial tuning curves

Tuning curves were computed by dividing the Gaussian-smoothed (σ = 3) number of significant calcium events in each of 100 spatial bins by the corresponding run-epoch occupancy. For each trial type, curves were normalized to the neuron’s peak activity.

###### Place field width

Place field width was determined from each neuron’s rate map, calculated as the run-epoch event count divided by occupancy across 100 spatial bins and smoothed with a Gaussian kernel (σ = 3). Local maxima were identified and fit with Gaussian functions, and field width was defined as the distance between positions at which the fitted curve reached 20% of its peak amplitude. Overlapping candidate fields were combined. Only fields containing at least five significant events on separate laps were analyzed.

###### Recurrence

Recurrence was calculated as the fraction of tuned cells that remain significantly tuned the following day.

#### Task-selective neurons

Tuned neurons were classified as A-selective if they were tuned only in A trials and not B, B-selective if they were tuned in B trials and not A, and AB-Shared if they were tuned in both A and B trials. As previously described^22^, neurons were included in the analysis if they exhibited at least five calcium transients within their place field during run epochs, with these events occurring on separate laps. In addition, for the corresponding spatial bins on other laps, the animal had to be engaged in running behavior for at least 80% of the time spanning the same spatial range as the identified calcium events.

##### Stability

###### PV correlation

Spatial firing rate maps (100 spatial bins) were computed for each neuron by dividing calcium event rate by occupancy. For each session and trial type, the rate maps of all neurons were assembled into a 2D matrix in which rows corresponded to neurons and columns to spatial bins. Thus, each column represented the population activity vector across all neurons at a given spatial location. To measure stability of all cells, population vectors from corresponding spatial bins were Pearson-correlated between trial types or imaging sessions. The mean correlation across all spatial bins was taken as the population vector (PV) correlation score. For off-diagonal quantification, all possible non-corresponding spatial bin pairs are compared between imaging sessions. PV correlation was calculated using all matched cells (main figure) and using only matched, tuned cells (supplement).

###### TC correlation

For spatially tuned neurons matched between two sessions, the spatial tuning vectors across 100 spatial bins were Pearson correlated for each neuron and the mean across all neurons was the tuning vector correlation score.

###### Centroid shift

Place field centroids were defined as the angle of each neuron’s maximal tuning vector (i.e., the place field with the highest event rate), represented in circular space as a complex vector. For each neuron, centroid shift was calculated as the minimal angular difference between centroids across sessions, constrained to the range 0–π radians to account for circular wrapping.

##### Decoding

###### PV decoding

A decoder was constructed for each session for each mouse as described previously^22^. Template tuning curves for each cell were constructed in a similar manner as described above, only using data from run epochs. Data from the first half of the session was used to define the template.

Time-resolved firing rate vectors were generated for each neuron from the second half of the session using 250 ms time bins and Gaussian smoothing (σ = 250 ms). At each time point, position was decoded by identifying the template location with the highest correlation to the population activity vector.

Trial classification error was calculated as the fraction of time points misassigned to A or B trials. Position decoding error was computed as the mean absolute difference between decoded and true position, collapsing across trial types. Distances were treated circularly, with positions 0 and 200 considered equivalent.

###### Statistics

All results are reported as mean ± SEM. Statistics were done in both GraphPad Prism and MATLAB. Normality was assessed using the Kolmogorov-Smirnov test to choose between parametric and non-parametric tests. All data points for both RF vs. OCGOL and Control vs. CNO are paired data within the same mouse. Statistical significance was assessed using the paired Student’s t test, paired Wilcoxon signed-rank test, repeated-measures one way ANOVA, Friedman test, Tukey post hoc test, Dunn post hoc test, and two-sided Pearson correlation where appropriate. The symbols *, **, and *** denote p values of <0.05, <0.01, and <0.001, respectively.

###### Author contributions

M.D.H and J.B. conceived the project and designed experiments; M.D.H performed all experiments and analysis for Figures 1-6. P.R. imaged all brain sections for histology and assisted in manual screening of ROIs. V.R. performed injections for a subset (n=4) of mice included for LEC silencing. E.K. processed and aligned histology imaged to a brain atlas. M.D.H generated all figures with input from J.B. M.D.H and J.B. wrote the manuscript.

## Acknowledgements

This work was supported by the following grants to J.B.: National Institutes of Health (NIH) National Institute of Neurological Disorders and Stroke (NINDS) BRAIN Initiative R01NS109994, NIH NINDS R01NS109362, NIH NINDS RM1NS132981, NIH National Institute of Mental Health (NIMH) R01MH122391, NIH National Institute on Aging (NIA) R01AG094086; Alzheimer’s Association AARGD-NTF-23-1151101, McKnight Scholar Award in Neuroscience, Klingenstein-Simons Fellowship Award in Neuroscience, Alfred P. Sloan Research Fellowship, Mathers Charitable Foundation Award, Whitehall Research Grant, a Blas Frangione Young Investigator Research Grant, New York University Whitehead Fellowship for Junior Faculty in Biomedical and Biological Sciences, and Leon Levy Foundation Award. M.D.H was supported by NIH 5T32MH096331-10, NIH 5T32NS086750-07, and NRSA 5TL1TR001447-09. V.R. was supported by a Young Researchers Bettencourt Prize. E.K. was supported by the R25 BRAIN Program 5R25NS125593 and NYU Dean’s Undergraduate Research Fund (DURF).

The New York University (NYU) High Performance Computing Cluster was used for data analysis. The NYU Genotyping Core Laboratory was used for genotyping mice.

We thank R. Zemla for the head-fixed behavioral setup and analysis pipeline. We thanks J.J. Moore for guidance on population vector decoding, feedback on analysis, and revision of the manuscript. We are grateful to G. Buzsáki, C. Constantinople, E. Klann, K. ONeil, M. Hernández-Frausto, and S. Wang for their input throughout the project and revisions of the manuscript.

## Materials & correspondence

Material requests and correspondence to Jayeeta Basu

## Data availability

Data will be available upon request to Jayeeta Basu and custom code will be available at https://github.com/basulab-nyu/Hopkins_2026

