## Supplemental Figures and Tables for "Lateral entorhinal cortex supports behaviorally-induced hippocampal ensemble stability for reliable memory recall"

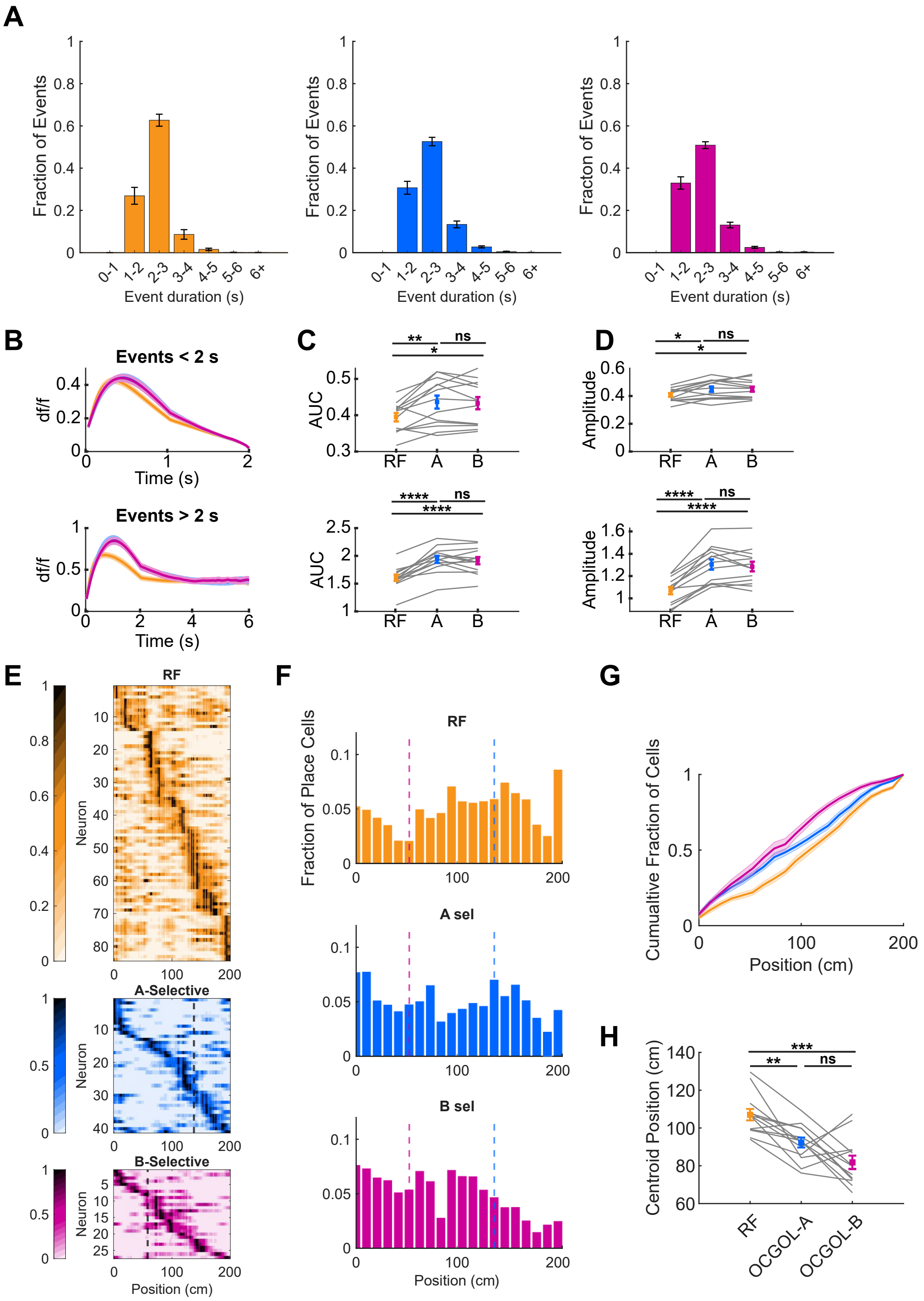
A
Fraction of Events
Event duration (s)
0-1
1-2
2-3
3-4
4-5
5-6
6+
Fracton of Events
B
Events < 2 s
Events > 2 s
df/f
Time (s)
C
AUC
RF
A
B
ns
D
Amplitude
E
RF
A-Selective
B-Selective
Neuron
Position (cm)
F
Fraction of Place Cells
A sel
B sel
G
Cumualtive Fraction of Cells
H
Centroid Position (cm)
OCGOL-A
OCGOL-B

**S1. Supplement to Figure 2**

A) Average histogram of duration of calcium events binned by 1 second for each task (RF- orange, OCGOL-A-blue, OCGOL-B-magenta). Each value corresponds to the average duration across all detected events for each mouse averaged across all mice.

B) ΔF/F trace of calcium events during run epochs (across all tasks) for events less than or equal to 2 seconds (top) and events greater than 2 seconds (bottom). Each value corresponds to the average ΔF/F trace across all detected events for each mouse averaged across all mice.

C) Area under the curve (AUC) of detected calcium events (across all tasks) during run epochs parsed by short duration events (top) and long duration events (bottom). Each value corresponds to the average AUC across all detected events for each mouse averaged across all mice. (Short duration events (top): one way RM ANOVA, $p=0.0021$; Tukey *post-hoc* test: RF vs. OCGOL-A, $p=0.0091$; RF vs. OCGOL-B, $p=0.0171$; OCGOL-A vs. OCGOL-B, $p=0.9020$. Long duration events (bottom): one way RM ANOVA, $p<0.0001$; Tukey *post-hoc* test: RF vs. OCGOL-A, $p<0.0001$; RF vs. OCGOL-B, $p<0.0001$; OCGOL-A vs OCGOL-B, $p=0.7750$).

D) Amplitude of detected calcium events during run epochs (across all tasks) parsed by short duration events (top) and long duration events (bottom). Each value corresponds to the average amplitude across all detected events for each mouse averaged across all mice (Short duration events (top): one way RM ANOVA, $p=0.0042$; Tukey *post-hoc* test: RF vs. OCGOL-A, $p=0.0169$; RF vs. OCGOL-B, $p=0.0271$; OCGOL-A vs. OCGOL-B, $p>0.9999$. Long duration events (bottom): one way RM ANOVA, $p<0.0001$; Tukey *post-hoc* test: RF vs. OCGOL-A, $p<0.0001$; RF vs. OCGOL-B, $p<0.0001$; OCGOL-A vs. OCGOL-B, $p=0.6343$).

E) Example rate map from a single mouse for cells tuned in RF (orange), cells selectively tuned in A trials (blue), and cells selectively tuned in B trials (magenta). Black dashed lines represent respective, fixed reward zones.

F) Distribution of tuned (RF) or trial-selective (OCGOL) place field centroids per 10-cm spatial bin across RF. Dashed lines represent A (blue) and B (magenta) reward zones.

G) Cumulative distribution of place field centroids across the track.

H) Mean place field centroid for tuned cells (RF) or trial-selective cells (OCGOL) (one way RM ANOVA, $p<0.0001$; Tukey *post-hoc* test: RF vs. OCGOL-A, $p=0.0019$; RF vs. OCGOL-B, $p=0.0003$, OCGOL-A vs. OCGOL-B, $p=0.1163$).

All data presented in figure: $n=13$ mice unless otherwise specified, means plotted with SEM.

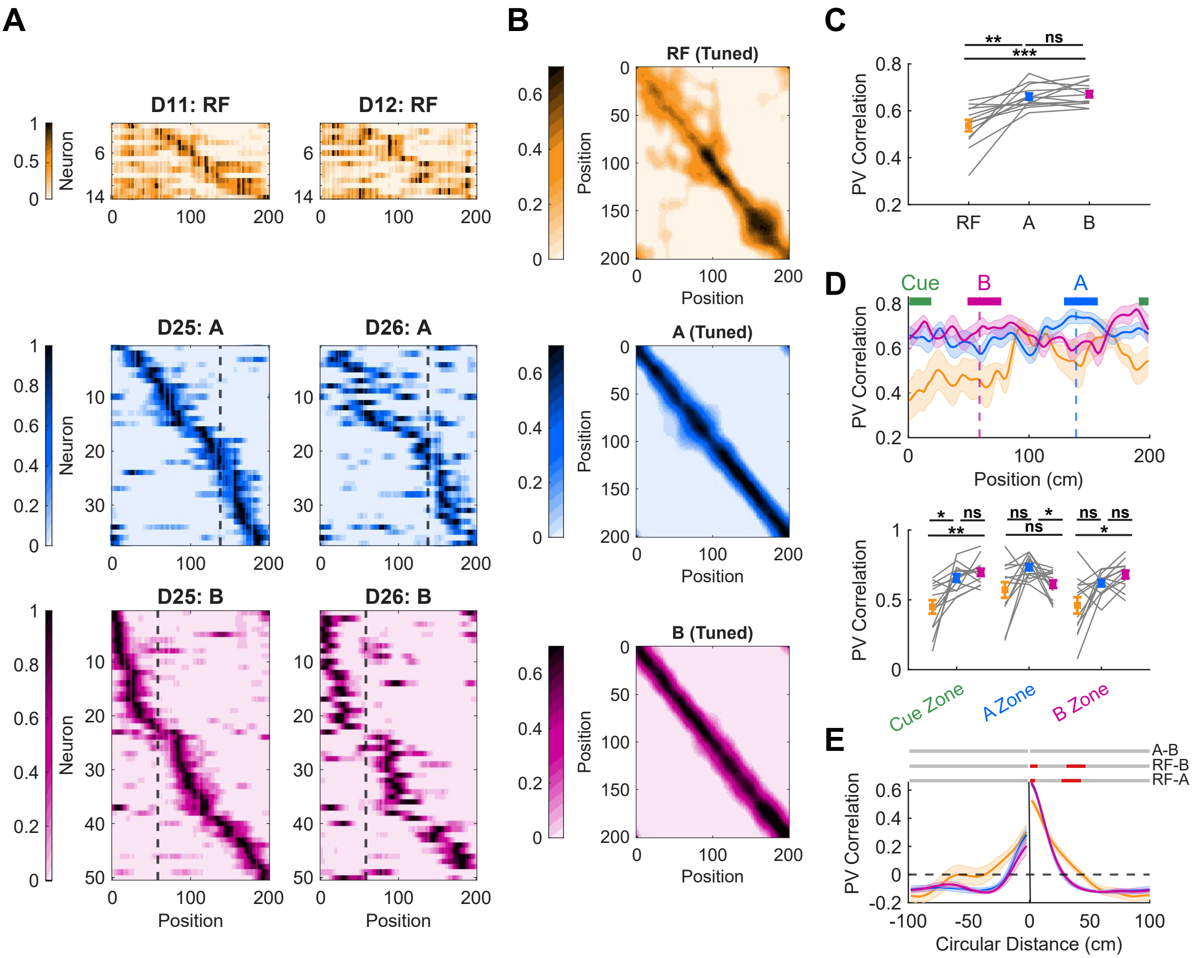

A
D11: RF
D12: RF
D25: A
D26: A
D25: B
D26: B
Neuron
Position
B
RF (Tuned)
A (Tuned)
B (Tuned)
Position
C
PV Correlation
RF
A
B
**
ns
***
D
Cue
B
A
PV Correlation
Position (cm)
Cue Zone
A Zone
B Zone
E
A-B
RF-B
RF-A
PV Correlation
Circular Distance (cm)

**S2. Supplement to Figure 3**

A) Example rate maps from a single mouse for all tuned and matched cells on day 1 (left) and day 2 (right) for each task. Each neuron's rate is normalized to its maximum rate and sorted according to the position of its maximum rate across 100 spatial bins.
B) Mean PV correlation maps for tuned cells across all mice for each task.
C) Mean PV correlation of tuned cells for neighboring days (one way RM ANOVA, $p<0.0001$; Tukey *post-hoc* test: RF vs. OCGOL-A, $p=0.0018$; RF vs. OCGOL-B, $p=0.0004$; OCGOL-A vs. OCGOL-B, $p=0.725$).
D) PV correlation of tuned cells plotted by spatial bin for each task (top). See Summary table for significant bin-wise p-values. Mean PV correlations of tuned cells for 30 cm zones centered at salient sensory locations (bottom): odor presentation (green), reward zone B (magenta), and reward zone A (blue) (Cue Zone: one way RM ANOVA, $p=0.0007$; Tukey post-hoc and paired t-test: RF vs. OCGOL-A, $p=0.0128$, $p=0.0050$ (respectively); RF vs OCGOL-B, $p=0.0017$, $p=0.0006$; OCGOL-A vs. OCGOL-B, $p=0.3751$, $p=0.1889$. Zone A: one way RM ANOVA, $p=0.0283$; Tukey *post-hoc* and paired t-test: RF vs. OCGOL-A, $p=0.0626$, $p=0.0260$ (respectively); RF vs OCGOL-B, $p=0.7638$, $p=0.4928$; OCGOL-A vs. OCGOL-B, $p=0.0273$, $p=0.0109$. Zone B: one way RM ANOVA, $p=0.0085$; Tukey *post-hoc* and paired t-test: RF vs. OCGOL-A, $p=0.0707$, $p=0.0295$ (respectively); RF vs OCGOL-B, $p=0.0189$, $p=0.0075$; OCGOL-A vs. OCGOL-B, $p=0.1221$, $p=0.0531$).
E) PV correlation for tuned cells of opposing spatial bins using off-diagonal values for each task (Friedman test, Effect of task $p=0.0079$; paired Wilcoxon signed-rank *post-hoc* test with FDR correction: RF vs. OCGOL-A: $p=0.0861$, RF vs. OCGOL-B: $p=0.0861$, OCGOL-A vs. OCGOL-B: $p=0.5417$). Horizontal bars represent p-values for all pairwise task combinations for a given spatial bin by paired Wilcoxon signed rank test (gray: $p>0.05$, red: $p<=0.05$). See Summary table for significant bin-wise p-values.

All data presented in figure: n=13 mice unless otherwise specified, means plotted with SEM.

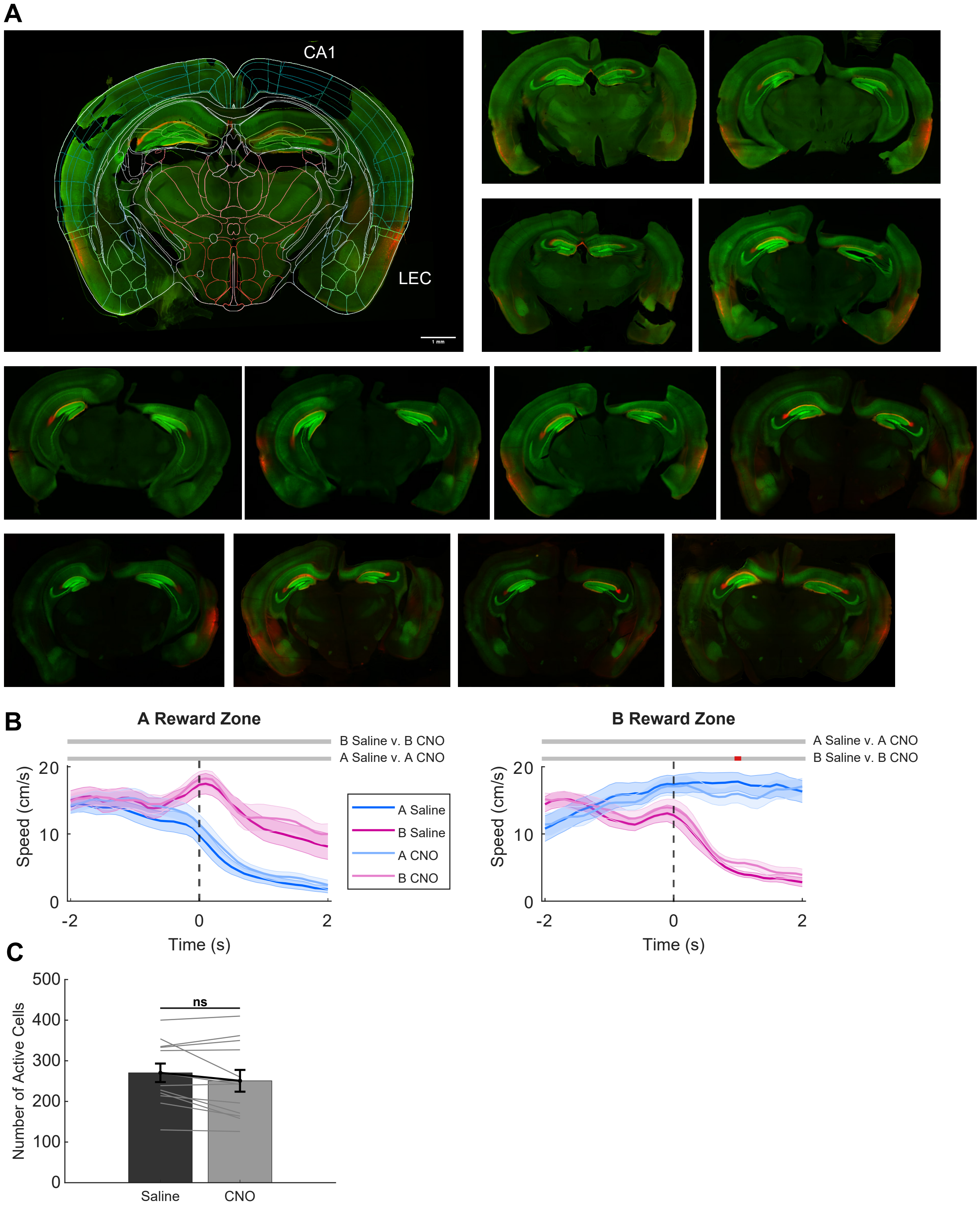

A
CA1
LEC
1 mm
B
A Reward Zone
B Saline v. B CNO
A Saline v. A CNO
Speed (cm/s)
Time (s)
A Saline
B Saline
A CNO
B CNO
B Reward Zone
A Saline v. A CNO
B Saline v. B CNO
C
Number of Active Cells
ns
Saline
CNO

**S3. Supplement to Figure 4**

A) Epifluorescent images of representative viral injection of AAV2.5-CaMKIIa-hM4D(Gi)-mCherry in LEC for each mouse with overlaid Allen Brain Atlas (n=13).

B) Mean animal speed (cm/s) 2 seconds before and after entry to A reward zone (left) and B reward zone (right) denoted by black dashed line. Color of line corresponds to the trial type (blue-A, magenta-B) and darkness corresponds to treatment (dark-saline, light-CNO) (n=13, Friedman test, A reward zone: A-Saline vs. CNO, Effect of task p=0.0803; A reward zone: B-Saline vs. CNO, Effect of treatment p=0.3396; B reward zone: B-Saline vs. CNO, Effect of treatment p=0.1272; B reward zone: A Saline vs. CNO, Effect of treatment p=0.1465. See Summary table for significant bin-wise p-values.

C) Mean number of active cells on day 2 saline (dark gray) vs. day 3 CNO (light gray) (n=12, Paired t-test, p=0.0893).

All data presented in figure: means plotted with SEM.

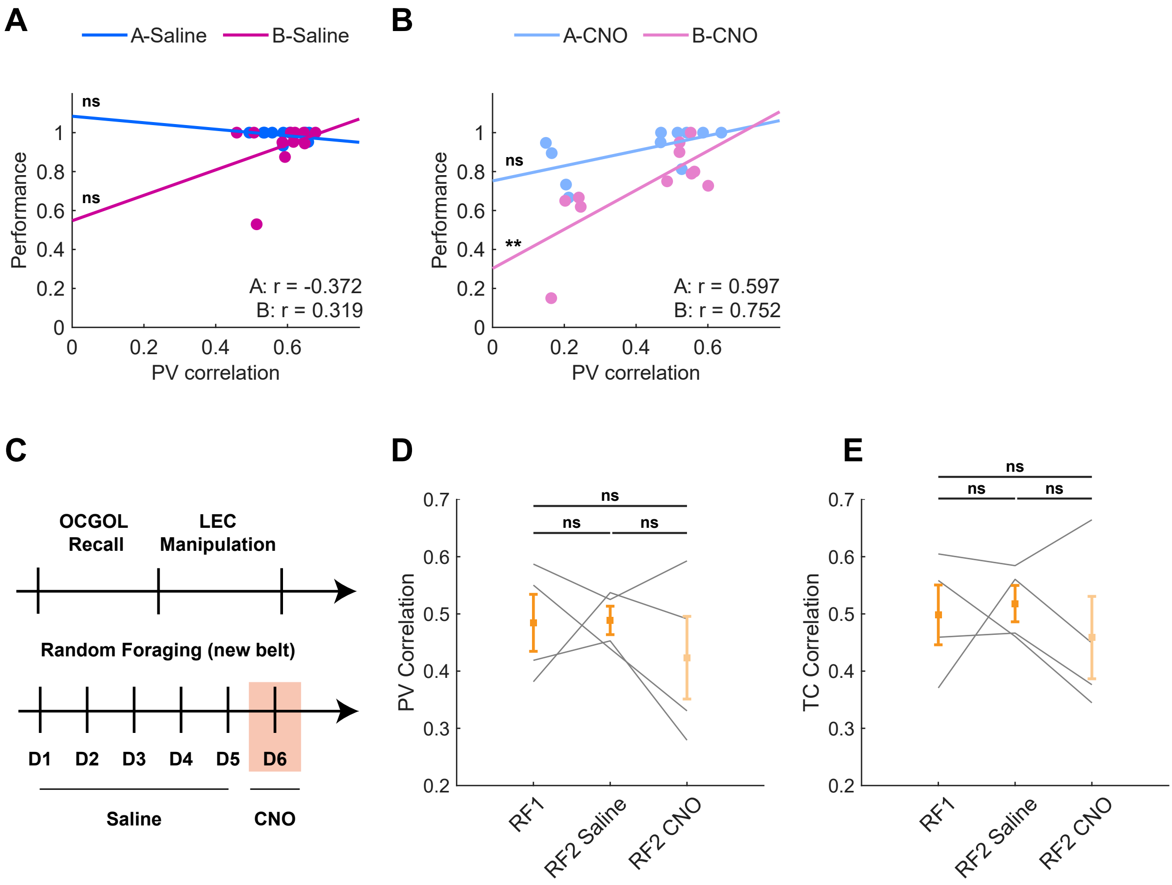

A
A-Saline
B-Saline
ns
ns
Performance
PV correlation
A: r = -0.372
B: r = 0.319
B
A-CNO
B-CNO
ns
**
A: r = 0.597
B: r = 0.752
C
OCGOL Recall
LEC Manipulation
Random Foraging (new belt)
D1
D2
D3
D4
D5
D6
Saline
CNO
D
PV Correlation
ns
RF1
RF2 Saline
RF2 CNO
E
TC Correlation

**S4. Supplement to Fig 5**

A) PV correlation (day 1 saline vs. day 2 saline) is plotted against behavioral performance (day 2 saline) during A (blue) and B trials (magenta) (n=11, Two-sided Pearson correlation, A: r=-0.372, p=0.260; B: r=0.319, p=0.340).
B) PV correlation (day 2 saline vs. day 3 CNO) is plotted against behavioral performance (day 3 CNO) (n=11, Two-sided Pearson correlation, A: r=0.597, p=0.052; B: r=0.752, p=0.008).
C) Behavioral timeline of RF sessions that occur after OCGOL and manipulation.
D) PV correlation of active CA1 neurons across RF1, RF2 Saline, and RF2 CNO (n=4, paired Wilcoxon signed rank test: RF1 vs. RF2 Saline, p>0.99; RF1 vs. RF2 CNO, p=0.625; RF2 Saline vs. RF2 CNO, p=0.3750).
E) TC correlation of tuned cells across all tasks (n=4, paired Wilcoxon signed rank test: RF1 vs. RF2 Saline, p>0.9999; RF1 vs. RF2 CNO, p=0.6250; RF2 Saline vs. RF2 CNO, p=0.2500).

All data presented in figure: means plotted with SEM.

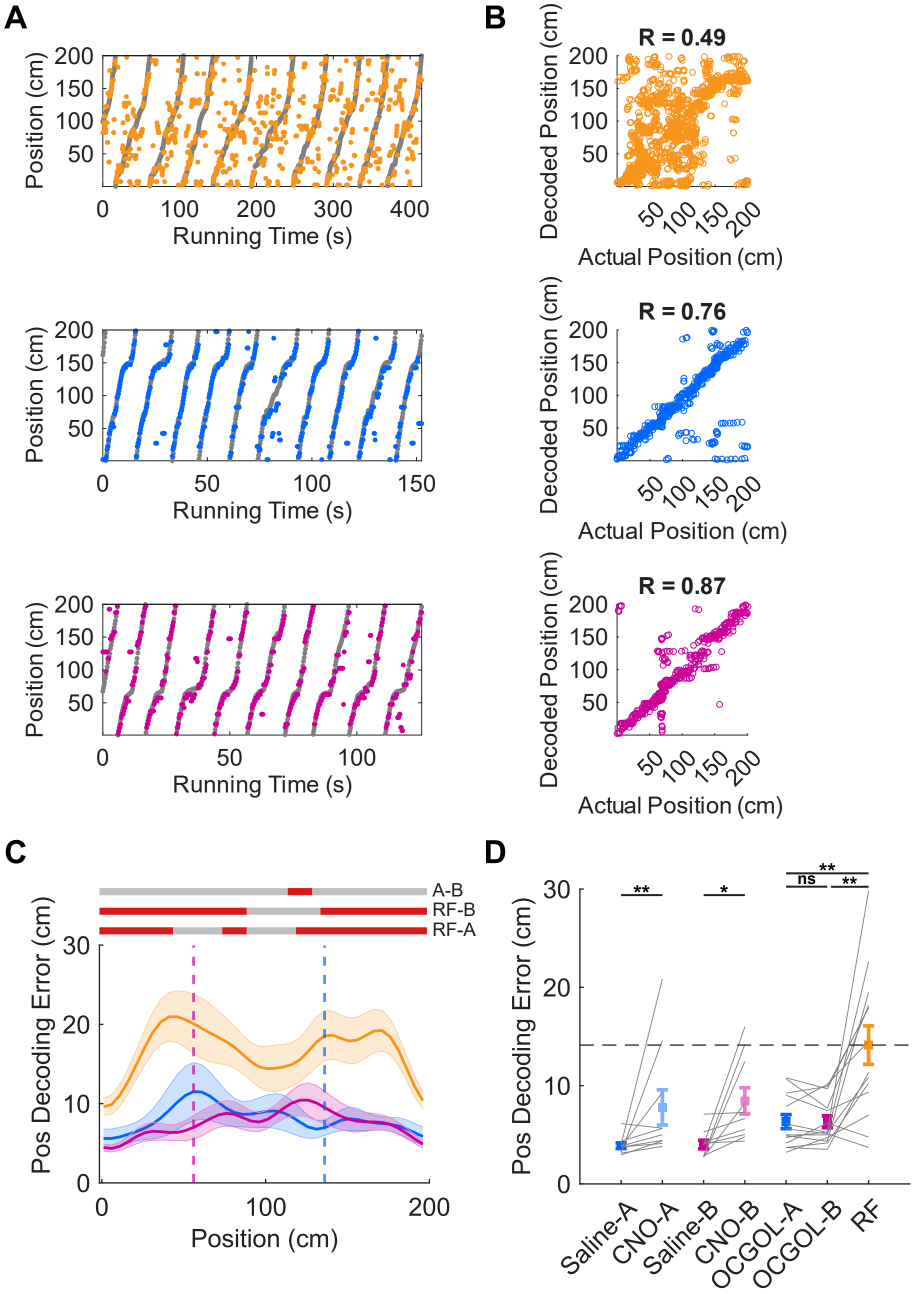
A
B
C
D
R = 0.49
R = 0.76
R = 0.87
Position (cm)
Running Time (s)
Decoded Position (cm)
Actual Position (cm)
A-B
RF-B
RF-A
Pos Decoding Error (cm)
Position (cm)
ns
Saline-A
CNO-A
Saline-B
CNO-B
OCGOL-A
OCGOL-B
RF

**S5. Supplement to Fig 6**

A) Example position decoding for a mouse across RF (top, orange), OCGOL-A (middle, blue), OCGOL-B (bottom, magenta) with the number of cells used to train the decoder equal amongst all tasks. Decoded positions are plotted on top of actual positions (gray) as a function of time.

B) Actual position plotted against decoded position for an example mouse across all tasks.

C) Position decoding error plotted by spatial bin for RF and OCGOL. Horizontal bars represent p-values for all pairwise task combinations for a given spatial bin by paired Wilcoxon signed rank test (gray: p>0.05, red: p<=0.05) (n=14). See Summary table for significant bin-wise p-values.

D) Position decoding error for animals that underwent LEC silencing (left four columns) and those that completed RF and OCGOL (right three columns). For RF vs. OCGOL comparisons, number of cells used to train the PV decoder were kept equal for each task per mouse. Dashed horizontal gray line at the mean position decoding error during RF (A-Saline vs. CNO: n=10, Wilcoxon signed rank test, p=0.0059; B-Saline vs. CNO, Paired t-test, p=0.0168; RF vs. OCGOL comparison: n=14, Friedman test, p=0.0015; Dunn's multiple comparisons *post-hoc* test: RF vs. OCGOL-A, p=0.0075; RF vs OCGOL-B, p=0.0039; OCGOL-A vs. OCGOL-B, p>0.999)

All data presented in figure: means plotted with SEM.

### Data Tables and Supplemental Tables

#### Fig 2J. SI score by spatial bins (20 spatial bins)

**RF vs. OCGOL-A**

| bin | p | p_fdr |
|---|---|---|
| 1 | 0.000488 | 0.000751 |
| 2 | 0.000977 | 0.001149 |
| 3 | 0.000244 | 0.000751 |
| 4 | 0.000244 | 0.000751 |
| 5 | 0.001953 | 0.002056 |
| 6 | 0.000977 | 0.001149 |
| 7 | 0.000488 | 0.000751 |
| 8 | 0.000488 | 0.000751 |
| 9 | 0.000488 | 0.000751 |
| 10 | 0.000977 | 0.001149 |
| 11 | 0.000488 | 0.000751 |
| 12 | 0.000488 | 0.000751 |
| 13 | 0.000488 | 0.000751 |
| 14 | 0.000244 | 0.000751 |
| 15 | 0.001465 | 0.001628 |
| 16 | 0.000488 | 0.000751 |
| 17 | 0.000244 | 0.000751 |
| 18 | 0.004883 | 0.004883 |
| 19 | 0.000977 | 0.001149 |
| 20 | 0.000488 | 0.000751 |

**RF vs. OCGOL-B**

| bin | p | p_fdr |
|---|---|---|
| 1 | 0.000977 | 0.002663 |
| 2 | 0.001953 | 0.003005 |
| 3 | 0.000244 | 0.002441 |
| 4 | 0.000244 | 0.002441 |
| 5 | 0.003906 | 0.004596 |
| 6 | 0.001953 | 0.003005 |
| 7 | 0.000488 | 0.002441 |
| 8 | 0.000977 | 0.002663 |
| 9 | 0.002441 | 0.003255 |
| 10 | 0.001465 | 0.002663 |
| 11 | 0.001465 | 0.002663 |
| 12 | 0.000488 | 0.002441 |
| 13 | 0.001221 | 0.002663 |
| 14 | 0.001221 | 0.002663 |
| 15 | 0.024414 | 0.024414 |
| 16 | 0.002441 | 0.003255 |
| 17 | 0.003418 | 0.004272 |
| 18 | 0.009766 | 0.010280 |
| 19 | 0.006836 | 0.007595 |
| 20 | 0.001465 | 0.002663 |

#### Fig 3G. PV by spatial bin (100 spatial bins)

**RF vs. OCGOL-A**

| bin | p | p_fdr |
|---|---|---|
| 1 | 0.008056641 | 0.040965969 |
| 43 | 0.006103516 | 0.034548202 |
| 63 | 0.006103516 | 0.034548202 |
| 64 | 0.003417969 | 0.025634766 |
| 65 | 0.000732422 | 0.010986328 |
| 66 | 0.000244141 | 0.009155273 |
| 67 | 0.001220703 | 0.014648438 |
| 68 | 0.004638672 | 0.028991699 |
| 79 | 0.010498047 | 0.048452524 |
| 80 | 0.010498047 | 0.048452524 |
| 81 | 0.008056641 | 0.040965969 |
| 82 | 0.003417969 | 0.025634766 |
| 83 | 0.001708984 | 0.017679149 |
| 84 | 0.002441406 | 0.020926339 |

**RF vs. OCGOL-B**

| bin | p | p_fdr |
|---|---|---|
| 1 | 0.002441 | 0.020926 |
| 2 | 0.004639 | 0.028992 |
| 3 | 0.010498 | 0.048453 |
| 8 | 0.003418 | 0.025635 |
| 9 | 0.002441 | 0.020926 |
| 10 | 0.003418 | 0.025635 |
| 11 | 0.004639 | 0.028992 |
| 16 | 0.008057 | 0.040966 |
| 17 | 0.006104 | 0.034548 |
| 18 | 0.002441 | 0.020926 |
| 19 | 0.001709 | 0.017679 |
| 20 | 0.004639 | 0.028992 |
| 21 | 0.008057 | 0.040966 |
| 33 | 0.008057 | 0.040966 |

**OCGOL-A vs. B**

| bin | p | p_fdr |
|---|---|---|
| 75 | 0.006104 | 0.034548 |
| 76 | 0.004639 | 0.028992 |
| 77 | 0.002441 | 0.020926 |
| 78 | 0.004639 | 0.028992 |
| 79 | 0.010498 | 0.048453 |

| | | |
|---|---|---|
| 85 | 0.001708984 | 0.017679149 |
| 86 | 0.000488281 | 0.010986328 |
| 87 | 0.000488281 | 0.010986328 |
| 88 | 0.000488281 | 0.010986328 |
| 89 | 0.000488281 | 0.010986328 |
| 90 | 0.000244141 | 0.009155273 |
| 91 | 0.000244141 | 0.009155273 |
| 92 | 0.000732422 | 0.010986328 |
| 93 | 0.001220703 | 0.014648438 |
| 94 | 0.000732422 | 0.010986328 |
| 95 | 0.000488281 | 0.010986328 |
| 96 | 0.003417969 | 0.025634766 |

| | | |
|---|---|---|
| 34 | 0.001709 | 0.017679 |
| 35 | 0.000732 | 0.010986 |
| 36 | 0.004639 | 0.028992 |
| 43 | 0.010498 | 0.048453 |
| 44 | 0.006104 | 0.034548 |
| 85 | 0.008057 | 0.040966 |
| 86 | 0.001221 | 0.014648 |
| 87 | 0.000244 | 0.009155 |
| 88 | 0.000244 | 0.009155 |
| 89 | 0.000732 | 0.010986 |
| 90 | 0.000732 | 0.010986 |
| 91 | 0.001221 | 0.014648 |
| 92 | 0.001221 | 0.014648 |
| 93 | 0.000732 | 0.010986 |
| 94 | 0.000244 | 0.009155 |
| 95 | 0.000244 | 0.009155 |
| 96 | 0.000244 | 0.009155 |
| 97 | 0.002441 | 0.020926 |
| 98 | 0.004639 | 0.028992 |
| 99 | 0.010498 | 0.048453 |

**Fig 3H. PV off-diagonal by bin difference (99 spatial bins)**

**RF vs. OCGOL-A**

| bin | p | p_fdr | bin | p | p_fdr |
|---|---|---|---|---|---|
| 1 | 0.004638672 | 0.009840611 | 50 | 0.002441406 | 0.006713867 |
| 2 | 0.006103516 | 0.012416056 | 51 | 0.004638672 | 0.009840611 |
| 3 | 0.004638672 | 0.009840611 | 52 | 0.013427734 | 0.023880462 |
| 4 | 0.004638672 | 0.009840611 | 61 | 0.013427734 | 0.023880462 |
| 5 | 0.006103516 | 0.012416056 | 62 | 0.010498047 | 0.019487 |
| 6 | 0.002441406 | 0.006713867 | 63 | 0.004638672 | 0.009840611 |
| 7 | 0.002441406 | 0.006713867 | 64 | 0.004638672 | 0.009840611 |
| 8 | 0.004638672 | 0.009840611 | 65 | 0.002441406 | 0.006713867 |
| 9 | 0.003417969 | 0.00838956 | 66 | 0.001708984 | 0.005767822 |
| 10 | 0.002441406 | 0.006713867 | 67 | 0.001220703 | 0.0053316 |
| 11 | 0.002441406 | 0.006713867 | 68 | 0.001220703 | 0.0053316 |
| 12 | 0.002441406 | 0.006713867 | 69 | 0.001220703 | 0.0053316 |
| 13 | 0.002441406 | 0.006713867 | 70 | 0.001220703 | 0.0053316 |
| 14 | 0.001708984 | 0.005767822 | 71 | 0.000732422 | 0.005305593 |
| 15 | 0.001708984 | 0.005767822 | 72 | 0.001220703 | 0.0053316 |
| 16 | 0.001708984 | 0.005767822 | 73 | 0.001220703 | 0.0053316 |

| | | | | | |
|---|---|---|---|---|---|
| 17 | 0.001708984 | 0.005767822 | 74 | 0.001220703 | 0.0053316 |
| 18 | 0.001708984 | 0.005767822 | 75 | 0.001220703 | 0.0053316 |
| 19 | 0.001220703 | 0.0053316 | 76 | 0.001220703 | 0.0053316 |
| 20 | 0.001220703 | 0.0053316 | 77 | 0.000732422 | 0.005305593 |
| 21 | 0.001220703 | 0.0053316 | 78 | 0.001220703 | 0.0053316 |
| 22 | 0.001220703 | 0.0053316 | 79 | 0.001220703 | 0.0053316 |
| 23 | 0.001220703 | 0.0053316 | 80 | 0.001708984 | 0.005767822 |
| 24 | 0.001220703 | 0.0053316 | 81 | 0.002441406 | 0.006713867 |
| 25 | 0.001220703 | 0.0053316 | 82 | 0.003417969 | 0.00838956 |
| 26 | 0.000732422 | 0.005305593 | 83 | 0.008056641 | 0.015846505 |
| 27 | 0.000488281 | 0.00426528 | 84 | 0.010498047 | 0.019487 |
| 28 | 0.000488281 | 0.00426528 | 85 | 0.010498047 | 0.019487 |
| 29 | 0.000488281 | 0.00426528 | 86 | 0.013427734 | 0.023880462 |
| 30 | 0.000244141 | 0.003816303 | 87 | 0.010498047 | 0.019487 |
| 31 | 0.000244141 | 0.003816303 | 88 | 0.008056641 | 0.015846505 |
| 32 | 0.000244141 | 0.003816303 | 89 | 0.013427734 | 0.023880462 |
| 33 | 0.000244141 | 0.003816303 | 90 | 0.013427734 | 0.023880462 |
| 34 | 0.000244141 | 0.003816303 | 91 | 0.010498047 | 0.019487 |
| 35 | 0.000244141 | 0.003816303 | 92 | 0.010498047 | 0.019487 |
| 36 | 0.000244141 | 0.003816303 | 93 | 0.008056641 | 0.015846505 |
| 37 | 0.000244141 | 0.003816303 | 94 | 0.003417969 | 0.00838956 |
| 38 | 0.000244141 | 0.003816303 | 95 | 0.002441406 | 0.006713867 |
| 39 | 0.000488281 | 0.00426528 | 96 | 0.003417969 | 0.00838956 |
| 40 | 0.000488281 | 0.00426528 | 97 | 0.003417969 | 0.00838956 |
| 41 | 0.000488281 | 0.00426528 | 98 | 0.002441406 | 0.006713867 |
| 42 | 0.000732422 | 0.005305593 | 99 | 0.003417969 | 0.00838956 |
| 43 | 0.002441406 | 0.006713867 | | | |

**RF vs. OCGOL-B**

| bin | p | p_fdr | bin | p | p_fdr |
|---|---|---|---|---|---|
| 1 | 0.001708984 | 0.005767822 | 50 | 0.000732422 | 0.005305593 |
| 2 | 0.001708984 | 0.005767822 | 51 | 0.002441406 | 0.006713867 |
| 3 | 0.001708984 | 0.005767822 | 52 | 0.006103516 | 0.012416056 |
| 4 | 0.010498047 | 0.019487 | 53 | 0.017089844 | 0.030212402 |
| 5 | 0.010498047 | 0.019487 | 62 | 0.013427734 | 0.023880462 |
| 6 | 0.010498047 | 0.019487 | 63 | 0.008056641 | 0.015846505 |
| 7 | 0.013427734 | 0.023880462 | 64 | 0.008056641 | 0.015846505 |
| 8 | 0.004638672 | 0.009840611 | 65 | 0.006103516 | 0.012416056 |
| 9 | 0.003417969 | 0.00838956 | 66 | 0.006103516 | 0.012416056 |
| 10 | 0.004638672 | 0.009840611 | 67 | 0.004638672 | 0.009840611 |
| 11 | 0.004638672 | 0.009840611 | 68 | 0.003417969 | 0.00838956 |

| | | | | | |
|---|---|---|---|---|---|
| 12 | 0.003417969 | 0.00838956 | 69 | 0.001220703 | 0.0053316 |
| 13 | 0.002441406 | 0.006713867 | 70 | 0.001220703 | 0.0053316 |
| 14 | 0.002441406 | 0.006713867 | 71 | 0.001708984 | 0.005767822 |
| 15 | 0.001708984 | 0.005767822 | 72 | 0.001708984 | 0.005767822 |
| 16 | 0.001708984 | 0.005767822 | 73 | 0.001708984 | 0.005767822 |
| 17 | 0.001220703 | 0.0053316 | 74 | 0.001220703 | 0.0053316 |
| 18 | 0.000732422 | 0.005305593 | 75 | 0.001220703 | 0.0053316 |
| 19 | 0.000732422 | 0.005305593 | 76 | 0.001220703 | 0.0053316 |
| 20 | 0.000488281 | 0.00426528 | 77 | 0.001220703 | 0.0053316 |
| 21 | 0.000244141 | 0.003816303 | 78 | 0.001220703 | 0.0053316 |
| 22 | 0.000244141 | 0.003816303 | 79 | 0.002441406 | 0.006713867 |
| 23 | 0.000488281 | 0.00426528 | 80 | 0.003417969 | 0.00838956 |
| 24 | 0.000488281 | 0.00426528 | 81 | 0.003417969 | 0.00838956 |
| 25 | 0.000488281 | 0.00426528 | 82 | 0.004638672 | 0.009840611 |
| 26 | 0.000488281 | 0.00426528 | 83 | 0.004638672 | 0.009840611 |
| 27 | 0.000488281 | 0.00426528 | 84 | 0.004638672 | 0.009840611 |
| 28 | 0.000488281 | 0.00426528 | 85 | 0.004638672 | 0.009840611 |
| 29 | 0.000488281 | 0.00426528 | 86 | 0.004638672 | 0.009840611 |
| 30 | 0.000244141 | 0.003816303 | 87 | 0.004638672 | 0.009840611 |
| 31 | 0.000244141 | 0.003816303 | 88 | 0.004638672 | 0.009840611 |
| 32 | 0.000244141 | 0.003816303 | 89 | 0.004638672 | 0.009840611 |
| 33 | 0.000244141 | 0.003816303 | 90 | 0.003417969 | 0.00838956 |
| 34 | 0.000244141 | 0.003816303 | 91 | 0.002441406 | 0.006713867 |
| 35 | 0.000244141 | 0.003816303 | 92 | 0.002441406 | 0.006713867 |
| 36 | 0.000244141 | 0.003816303 | 93 | 0.003417969 | 0.00838956 |
| 37 | 0.000244141 | 0.003816303 | 94 | 0.002441406 | 0.006713867 |
| 38 | 0.000488281 | 0.00426528 | 95 | 0.001708984 | 0.005767822 |
| 39 | 0.001220703 | 0.0053316 | 96 | 0.001708984 | 0.005767822 |
| 40 | 0.002441406 | 0.006713867 | 97 | 0.001708984 | 0.005767822 |
| 41 | 0.006103516 | 0.012416056 | 98 | 0.001708984 | 0.005767822 |
| 42 | 0.021484375 | 0.037756564 | 99 | 0.001708984 | 0.005767822 |

**5H. PV spatial bin (Saline vs. CNO) (100 spatial bins)**

**A Saline vs. A CNO**

| **bin** | **p** | **p_fdr** | **bin** | **p** | **p_fdr** |
|---|---|---|---|---|---|
| 1 | 0.013671875 | 0.032552083 | 55 | 0.013672 | 0.032552 |
| 2 | 0.013671875 | 0.032552083 | 56 | 0.004883 | 0.025699 |
| 3 | 0.006835938 | 0.02734375 | 57 | 0.004883 | 0.025699 |
| 4 | 0.004882813 | 0.025699013 | 58 | 0.004883 | 0.025699 |
| 5 | 0.006835938 | 0.02734375 | 59 | 0.024414 | 0.042832 |
| 6 | 0.009765625 | 0.031502016 | 66 | 0.018555 | 0.037109 |

| | | | | | |
|---|---|---|---|---|---|
| 7 | 0.018554688 | 0.037109375 | 67 | 0.013672 | 0.032552 |
| 8 | 0.024414063 | 0.042831689 | 68 | 0.018555 | 0.037109 |
| 9 | 0.024414063 | 0.042831689 | 70 | 0.024414 | 0.042832 |
| 10 | 0.013671875 | 0.032552083 | 71 | 0.018555 | 0.037109 |
| 11 | 0.006835938 | 0.02734375 | 72 | 0.024414 | 0.042832 |
| 12 | 0.006835938 | 0.02734375 | 73 | 0.024414 | 0.042832 |
| 13 | 0.002929688 | 0.025699013 | 74 | 0.009766 | 0.031502 |
| 14 | 0.004882813 | 0.025699013 | 75 | 0.004883 | 0.025699 |
| 15 | 0.004882813 | 0.025699013 | 76 | 0.001953 | 0.025699 |
| 16 | 0.009765625 | 0.031502016 | 77 | 0.000977 | 0.016276 |
| 17 | 0.018554688 | 0.037109375 | 78 | 0.00293 | 0.025699 |
| 24 | 0.024414063 | 0.042831689 | 79 | 0.018555 | 0.037109 |
| 25 | 0.013671875 | 0.032552083 | 87 | 0.009766 | 0.031502 |
| 26 | 0.009765625 | 0.031502016 | 88 | 0.009766 | 0.031502 |
| 27 | 0.006835938 | 0.02734375 | 89 | 0.013672 | 0.032552 |
| 28 | 0.002929688 | 0.025699013 | 90 | 0.013672 | 0.032552 |
| 29 | 0.002929688 | 0.025699013 | 95 | 0.018555 | 0.037109 |
| 30 | 0.004882813 | 0.025699013 | 96 | 0.000977 | 0.016276 |
| 31 | 0.006835938 | 0.02734375 | 97 | 0.000977 | 0.016276 |
| 32 | 0.013671875 | 0.032552083 | 98 | 0.000977 | 0.016276 |
| 44 | 0.013671875 | 0.032552083 | 99 | 0.000977 | 0.016276 |
| 45 | 0.013671875 | 0.032552083 | 100 | 0.000977 | 0.016276 |
| 46 | 0.018554688 | 0.037109375 | | | |

**B Saline vs. B CNO**

| bin | p | p_fdr | bin | p | p_fdr |
|---|---|---|---|---|---|
| 9 | 0.018554688 | 0.037866709 | 50 | 0.013672 | 0.031795 |
| 10 | 0.009765625 | 0.030517578 | 51 | 0.013672 | 0.031795 |
| 11 | 0.006835938 | 0.028483073 | 52 | 0.006836 | 0.028483 |
| 12 | 0.009765625 | 0.030517578 | 53 | 0.006836 | 0.028483 |
| 13 | 0.018554688 | 0.037866709 | 54 | 0.006836 | 0.028483 |
| 14 | 0.009765625 | 0.030517578 | 55 | 0.013672 | 0.031795 |
| 15 | 0.004882813 | 0.028483073 | 56 | 0.013672 | 0.031795 |
| 16 | 0.001953125 | 0.028483073 | 57 | 0.009766 | 0.030518 |
| 17 | 0.001953125 | 0.028483073 | 58 | 0.009766 | 0.030518 |
| 18 | 0.001953125 | 0.028483073 | 59 | 0.013672 | 0.031795 |
| 19 | 0.001953125 | 0.028483073 | 60 | 0.013672 | 0.031795 |
| 20 | 0.001953125 | 0.028483073 | 61 | 0.013672 | 0.031795 |
| 21 | 0.002929688 | 0.028483073 | 62 | 0.018555 | 0.037867 |
| 22 | 0.004882813 | 0.028483073 | 63 | 0.024414 | 0.046064 |
| 23 | 0.006835938 | 0.028483073 | 64 | 0.024414 | 0.046064 |

| | | | | | |
|---|---|---|---|---|---|
| 24 | 0.018554688 | 0.037866709 | 66 | 0.024414 | 0.046064 |
| 25 | 0.013671875 | 0.031795058 | 77 | 0.018555 | 0.037867 |
| 26 | 0.006835938 | 0.028483073 | 78 | 0.013672 | 0.031795 |
| 27 | 0.013671875 | 0.031795058 | 79 | 0.004883 | 0.028483 |
| 28 | 0.013671875 | 0.031795058 | 80 | 0.004883 | 0.028483 |
| 29 | 0.004882813 | 0.028483073 | 81 | 0.009766 | 0.030518 |
| 30 | 0.006835938 | 0.028483073 | 82 | 0.018555 | 0.037867 |
| 31 | 0.009765625 | 0.030517578 | 97 | 0.024414 | 0.046064 |
| 46 | 0.009765625 | 0.030517578 | 98 | 0.006836 | 0.028483 |
| 47 | 0.000976563 | 0.028483073 | 99 | 0.004883 | 0.028483 |
| 48 | 0.002929688 | 0.028483073 | 100 | 0.006836 | 0.028483 |
| 49 | 0.004882813 | 0.028483073 | | | |

**5K. PV correlation same day by spatial bin (100 spatial bins)**

**Saline vs. CNO**

| bin | p | p_fdr |
|---|---|---|
| 5 | 0.004882813 | 0.044389205 |
| 6 | 0.004882813 | 0.044389205 |
| 7 | 0.002929688 | 0.044389205 |
| 14 | 0.001953125 | 0.044389205 |
| 15 | 0.001953125 | 0.044389205 |
| 16 | 0.002929688 | 0.044389205 |
| 17 | 0.004882813 | 0.044389205 |
| 18 | 0.004882813 | 0.044389205 |
| 19 | 0.001953125 | 0.044389205 |
| 20 | 0.000976563 | 0.044389205 |
| 21 | 0.004882813 | 0.044389205 |

**6C. Trial identification error by spatial bin (40 spatial bins)**

**A Saline vs. A CNO**

| bin | p | p_fdr |
|---|---|---|
| 15 | 0.01171875 | 0.032169118 |
| 16 | 0.0078125 | 0.026041667 |
| 17 | 0.00390625 | 0.015625 |
| 18 | 0.00390625 | 0.015625 |
| 19 | 0.00390625 | 0.015625 |
| 20 | 0.0078125 | 0.026041667 |
| 21 | 0.013671875 | 0.032169118 |
| 22 | 0.013671875 | 0.032169118 |
| 23 | 0.013671875 | 0.032169118 |
| 24 | 0.013671875 | 0.032169118 |

**B Saline vs. B CNO**

| bin | p | p_fdr |
|---|---|---|
| 2 | 0.027344 | 0.047554 |
| 3 | 0.019531 | 0.045956 |
| 4 | 0.013672 | 0.036458 |
| 5 | 0.013672 | 0.036458 |
| 6 | 0.003906 | 0.021307 |
| 7 | 0.005859 | 0.021307 |
| 8 | 0.005859 | 0.021307 |
| 9 | 0.005859 | 0.021307 |
| 10 | 0.005859 | 0.021307 |
| 11 | 0.003906 | 0.021307 |

| | | |
|---|---|---|
| 25 | 0.001953125 | 0.015625 |
| 26 | 0.00390625 | 0.015625 |
| 27 | 0.00390625 | 0.015625 |
| 28 | 0.00390625 | 0.015625 |
| 29 | 0.00390625 | 0.015625 |
| 30 | 0.00390625 | 0.015625 |
| 31 | 0.00390625 | 0.015625 |

| | | |
|---|---|---|
| 12 | 0.003906 | 0.021307 |
| 13 | 0.003906 | 0.021307 |
| 14 | 0.009766 | 0.030048 |
| 15 | 0.027344 | 0.047554 |
| 25 | 0.027344 | 0.047554 |
| 26 | 0.027344 | 0.047554 |
| 27 | 0.027344 | 0.047554 |
| 28 | 0.019531 | 0.045956 |
| 29 | 0.009766 | 0.030048 |
| 30 | 0.005859 | 0.021307 |
| 31 | 0.005859 | 0.021307 |
| 32 | 0.005859 | 0.021307 |
| 33 | 0.027344 | 0.047554 |

**6E. Position decoding error by spatial bin (40 spatial bins)**

**A Saline vs. A CNO**

| | | |
|---|---|---|
| 16 | 0.005859375 | 0.026041667 |
| 17 | 0.005859375 | 0.026041667 |
| 18 | 0.001953125 | 0.015625 |
| 19 | 0.001953125 | 0.015625 |
| 20 | 0.001953125 | 0.015625 |
| 21 | 0.001953125 | 0.015625 |
| 22 | 0.001953125 | 0.015625 |
| 23 | 0.013671875 | 0.049715909 |
| 34 | 0.00390625 | 0.026041667 |
| 35 | 0.005859375 | 0.026041667 |

**6H. Position decoding error by spatial bin (40 spatial bins)**

**RF vs. OCGOL-A**

| bin | p | p_fdr |
|---|---|---|
| 1 | 0.000854492 | 0.009155273 |
| 2 | 0.002319336 | 0.009597252 |
| 3 | 0.00402832 | 0.011509487 |
| 4 | 0.008544922 | 0.021362305 |
| 5 | 0.010742188 | 0.024789663 |
| 6 | 0.010742188 | 0.024789663 |
| 7 | 0.010742188 | 0.024789663 |
| 8 | 0.013427734 | 0.02983941 |
| 9 | 0.016601563 | 0.03434806 |
| 10 | 0.016601563 | 0.03434806 |
| 11 | 0.020263672 | 0.040527344 |

**RF vs. OCGOL-B**

| bin | p | p_fdr |
|---|---|---|
| 1 | 0.001709 | 0.009322 |
| 2 | 0.001709 | 0.009322 |
| 3 | 0.004028 | 0.011509 |
| 4 | 0.005249 | 0.014316 |
| 5 | 0.004028 | 0.011509 |
| 6 | 0.005249 | 0.014316 |
| 7 | 0.004028 | 0.011509 |
| 8 | 0.003052 | 0.010771 |
| 9 | 0.001709 | 0.009322 |
| 10 | 0.000366 | 0.009155 |
| 11 | 0.000366 | 0.009155 |

| | | |
|---|---|---|
| 12 | 0.024536133 | 0.045297476 |
| 14 | 0.024536133 | 0.045297476 |
| 15 | 0.024536133 | 0.045297476 |
| 16 | 0.016601563 | 0.03434806 |
| 17 | 0.010742188 | 0.024789663 |
| 24 | 0.024536133 | 0.045297476 |
| 25 | 0.001708984 | 0.009321733 |
| 26 | 0.000610352 | 0.009155273 |
| 27 | 0.001220703 | 0.009155273 |
| 28 | 0.003051758 | 0.01077091 |
| 29 | 0.00402832 | 0.011509487 |
| 30 | 0.00402832 | 0.011509487 |
| 31 | 0.003051758 | 0.01077091 |
| 32 | 0.001708984 | 0.009321733 |
| 33 | 0.000854492 | 0.009155273 |
| 34 | 0.000610352 | 0.009155273 |
| 35 | 0.000854492 | 0.009155273 |
| 36 | 0.000854492 | 0.009155273 |
| 37 | 0.000854492 | 0.009155273 |
| 38 | 0.000854492 | 0.009155273 |
| 39 | 0.001220703 | 0.009155273 |
| 40 | 0.001220703 | 0.009155273 |

| | | |
|---|---|---|
| 12 | 0.000366 | 0.009155 |
| 13 | 0.001221 | 0.009155 |
| 14 | 0.004028 | 0.011509 |
| 15 | 0.004028 | 0.011509 |
| 16 | 0.008545 | 0.021362 |
| 17 | 0.006714 | 0.017514 |
| 18 | 0.016602 | 0.034348 |
| 19 | 0.024536 | 0.045297 |
| 28 | 0.020264 | 0.040527 |
| 29 | 0.013428 | 0.029839 |
| 30 | 0.006714 | 0.017514 |
| 31 | 0.003052 | 0.010771 |
| 32 | 0.003052 | 0.010771 |
| 33 | 0.002319 | 0.009597 |
| 34 | 0.002319 | 0.009597 |
| 35 | 0.001221 | 0.009155 |
| 36 | 0.001709 | 0.009322 |
| 37 | 0.002319 | 0.009597 |
| 38 | 0.002319 | 0.009597 |
| 39 | 0.002319 | 0.009597 |
| 40 | 0.002319 | 0.009597 |

**S2D. PV by spatial bin for tuned cells only (100 spatial bins)**

**RF vs. OCGOL-B**

| bin | p | p_fdr |
|---|---|---|
| 6 | 0.000732422 | 0.021972656 |
| 7 | 0.000488281 | 0.021972656 |
| 8 | 0.000732422 | 0.021972656 |
| 36 | 0.001708984 | 0.039438101 |
| 37 | 0.001708984 | 0.039438101 |
| 88 | 0.000244141 | 0.021972656 |
| 89 | 0.000244141 | 0.021972656 |
| 90 | 0.000732422 | 0.021972656 |
| 94 | 0.000732422 | 0.021972656 |
| 95 | 0.000488281 | 0.021972656 |
| 96 | 0.000732422 | 0.021972656 |

**OCGOL-A vs. B**

| bin | p | p_fdr |
|---|---|---|
| 94 | 0.000488281 | 0.021972656 |
| 95 | 0.001220703 | 0.033291903 |

**S2E. Off-diagonal by bin difference for tuned cells only (99 spatial bin distances)**

**RF vs. OCGOL-A**

| bin | p | p_fdr |
|---|---|---|
| 50 | 0.000244141 | 0.036254883 |
| 51 | 0.000488281 | 0.036254883 |
| 63 | 0.002441406 | 0.036254883 |
| 64 | 0.002441406 | 0.036254883 |
| 65 | 0.001220703 | 0.036254883 |
| 66 | 0.001220703 | 0.036254883 |
| 67 | 0.001708984 | 0.036254883 |
| 68 | 0.002441406 | 0.036254883 |
| 69 | 0.002441406 | 0.036254883 |
| 70 | 0.002441406 | 0.036254883 |

**RF vs. OCGOL-B**

| bin | p | p_fdr |
|---|---|---|
| 50 | 0.000244141 | 0.036254883 |
| 51 | 0.000488281 | 0.036254883 |
| 52 | 0.001220703 | 0.036254883 |
| 65 | 0.003417969 | 0.048339844 |
| 66 | 0.002441406 | 0.036254883 |
| 67 | 0.001708984 | 0.036254883 |
| 68 | 0.001708984 | 0.036254883 |
| 69 | 0.001708984 | 0.036254883 |
| 70 | 0.001708984 | 0.036254883 |
| 71 | 0.001708984 | 0.036254883 |
| 72 | 0.002441406 | 0.036254883 |

**S3B. Speed reward zone entry ± 2 seconds (Saline vs. CNO) (41 time bins centered at 0)**

**B Reward Zone for B Trials (Saline vs. CNO)**

| bin | p | p_fdr |
|---|---|---|
| 31 | 0.000732422 | 0.030029297 |

**S5C. Position decoding error by spatial bin (ROIs controlled per mouse) (40 spatial bins)**

**RF vs. OCGOL-A**

| bin | p | p_fdr |
|---|---|---|
| 1 | 0.001708984 | 0.00603171 |
| 2 | 0.005249023 | 0.014648438 |
| 3 | 0.002319336 | 0.007522171 |
| 4 | 0.003051758 | 0.009390024 |
| 5 | 0.00402832 | 0.012084961 |
| 6 | 0.005249023 | 0.014648438 |
| 7 | 0.008544922 | 0.020926339 |
| 8 | 0.010742188 | 0.024789663 |
| 9 | 0.024536133 | 0.047489289 |
| 16 | 0.013427734 | 0.029296875 |
| 17 | 0.006713867 | 0.018310547 |
| 18 | 0.020263672 | 0.040527344 |
| 25 | 0.008544922 | 0.020926339 |
| 26 | 0.001220703 | 0.004882813 |
| 27 | 0.000854492 | 0.004272461 |
| 28 | 0.000610352 | 0.00332919 |

**RF vs. OCGOL-B**

| bin | p | p_fdr |
|---|---|---|
| 1 | 0.000244 | 0.003329 |
| 2 | 0.000122 | 0.003329 |
| 3 | 0.000366 | 0.003329 |
| 4 | 0.00061 | 0.003329 |
| 5 | 0.001221 | 0.004883 |
| 6 | 0.001709 | 0.006032 |
| 7 | 0.001709 | 0.006032 |
| 8 | 0.001221 | 0.004883 |
| 9 | 0.00061 | 0.003329 |
| 10 | 0.000366 | 0.003329 |
| 11 | 0.000366 | 0.003329 |
| 12 | 0.001221 | 0.004883 |
| 13 | 0.020264 | 0.040527 |
| 14 | 0.020264 | 0.040527 |
| 15 | 0.010742 | 0.02479 |
| 16 | 0.008545 | 0.020926 |

**OCGOL-A vs. B**

| bin | p | p_fdr |
|---|---|---|
| 24 | 0.024536 | 0.047489 |
| 25 | 0.013428 | 0.029297 |
| 26 | 0.020264 | 0.040527 |

| | | |
|---|---|---|
| 29 | 0.000854492 | 0.004272461 |
| 30 | 0.001220703 | 0.004882813 |
| 31 | 0.000610352 | 0.00332919 |
| 32 | 0.000366211 | 0.00332919 |
| 33 | 0.000610352 | 0.00332919 |
| 34 | 0.000366211 | 0.00332919 |
| 35 | 0.000244141 | 0.00332919 |
| 36 | 0.000244141 | 0.00332919 |
| 37 | 0.000610352 | 0.00332919 |
| 38 | 0.001708984 | 0.00603171 |
| 39 | 0.003051758 | 0.009390024 |
| 40 | 0.001220703 | 0.004882813 |

| | | |
|---|---|---|
| 17 | 0.010742 | 0.02479 |
| 18 | 0.013428 | 0.029297 |
| 28 | 0.016602 | 0.035575 |
| 29 | 0.008545 | 0.020926 |
| 30 | 0.008545 | 0.020926 |
| 31 | 0.005249 | 0.014648 |
| 32 | 0.002319 | 0.007522 |
| 33 | 0.002319 | 0.007522 |
| 34 | 0.00061 | 0.003329 |
| 35 | 0.00061 | 0.003329 |
| 36 | 0.000366 | 0.003329 |
| 37 | 0.000366 | 0.003329 |
| 38 | 0.000366 | 0.003329 |
| 39 | 0.000366 | 0.003329 |
| 40 | 0.00061 | 0.003329 |
